# Structure-guided design of a CD81-binding mini-protein that blocks hepatitis C virus entry

**DOI:** 10.64898/2026.07.10.737702

**Authors:** Sojin An, Eunji Jo, Hee Seong Choi, Sang Soo Kim, Dong Won Kim, Woorim Yang, Sung Key Jang, Kyu-Ho Paul Park, Sang-Jun Ha, Yong Song Gho, Seong Wook Yang, Seungtaek Kim, Hyun-Soo Cho

**Affiliations:** Department of Systems Biology, College of Life Science and Biotechnology, Yonsei University, Seoul 03722, Republic of Korea; Applied Molecular Biochemistry Laboratory, Institut Pasteur Korea, Seongnam-si, Gyeonggi-do 13488, Republic of Korea; Department of Life Sciences, Pohang University of Science and Technology (POSTECH), Pohang 37673, Republic of Korea; Department of Biochemistry, College of Life Science and Biotechnology, Yonsei University, Seoul 03722, Republic of Korea; CEO Office, Institut Pasteur Korea, Seongnam-si, Gyeonggi-do 13488, Republic of Korea; Xenohelix Research Institute, BT Centre 305, 56 Songdogwahakro Yeonsugu, Incheon 21984, Republic of Korea; Zoonotic Virus Laboratory, Institut Pasteur Korea, Seongnam-si, Gyeonggi-do 13488, Republic of Korea

## Abstract

Despite the success of direct-acting antivirals, preventing hepatitis C virus (HCV) reinfection remains a critical global challenge. To address this, we leveraged deep learning-based *de novo* protein design to engineer mini-proteins targeting the large extracellular loop (LEL) of the HCV co-receptor CD81. These mini-proteins are predicted to precisely dock into CD81-LEL, occluding the critical binding interface required for the HCV E2 glycoprotein. Through biophysical screening, we identified mini-19, a lead candidate with sub-nanomolar affinity (*K*_D_ = 0.5 nM) and high thermostability up to 95°C. Flow cytometry and super-resolution imaging confirmed that mini-19 specifically recognizes the native topology of CD81 on cells and extracellular vesicles. Functionally, mini-19 neutralized HCVcc infection (IC_50_ = 1.2 nM). Molecular dynamics simulations demonstrated that mini-19 acts as a structural clamp, restricting the fusogenic conformational plasticity of the receptor. By targeting a conserved host factor rather than the rapidly mutating viral envelope, mini-19 provides a high genetic barrier to resistance. Beyond offering a prophylactic strategy against HCV, our findings establish a stable biologic platform for targeting tetraspanin-enriched microdomains and modulating host-pathogen interactions.

## Introduction

Hepatitis C virus (HCV) infection remains a global health burden despite the clinical success of direct-acting antivirals (DAAs) targeting the viral NS3/4A protease, NS5A, and NS5B polymerase (Kanwal *et al*, 2020; Lee *et al*, 2017; Smith *et al*, 2021). An estimated 70 million people worldwide are chronically infected. Hundreds of thousands of new cases continue to arise each year due to asymptomatic transmission, limited access to treatment, and high-risk exposure in vulnerable populations (Fasano *et al*, 2024; World Health Organization, 2025; Zhao *et al*, 2022b). Although DAAs achieve sustained virologic response rates exceeding 95%, their global impact is constrained by high costs, limited accessibility in low-and middle-income countries, and an inability to prevent new infections or reinfection (Rice, 2015; Shahid *et al*, 2021). This limitation leaves a significant prophylactic void. Because DAAs target intracellular replication, they do not block the earliest stages of the viral life cycle, including receptor-mediated entry, intercellular spread, or CD81-positive extracellular vesicle (EV)-facilitated transmission (Grakoui *et al*, 1993a, b). Consequently, HCV disseminates between hepatocytes, contributing to persistent infection, viral reseeding, and the burden of progressive cirrhosis, hepatocellular carcinoma, and liver transplantation (Bush *et al*, 2014; Cuypers *et al*, 2016; Das & Pandya, 2018; Shi *et al*, 2021; Zhao *et al*, 2022a).

These limitations highlight the need for complementary therapeutic strategies that target viral entry and the host factors essential for early infection. HCV utilizes multiple entry pathways, including receptor attachment, clathrin-dependent internalization, direct cell-to-cell transmission, and the uptake of CD81-positive EVs. These pathways operate independently of viral replication and remain unaffected by DAAs (Douam *et al*, 2015; Gerold *et al*, 2020). The tetraspanin CD81 is a central host factor required for HCV entry (Brazzoli *et al*, 2008; Higginbottom *et al*, 2000; Pileri *et al*, 1998). Its large extracellular loop (LEL), also known as extracellular loop 2, directly engages the HCV E2 glycoprotein. This interaction coordinates a receptor complex involving SR-BI, claudin-1, and occludin, enabling membrane fusion and productive entry (Evans *et al*, 2007; Harris *et al*, 2010; Harris *et al*, 2008; Kitadokoro *et al*, 2001; Kumar *et al*, 2021; Ploss *et al*, 2009; Scarselli *et al*, 2002). Beyond this role, CD81 organizes tetraspanin-enriched microdomains, modulates membrane fusion, and is enriched on hepatocyte-derived EVs that mediate viral dissemination (Hemler, 2005; Levy & Shoham, 2005; Masciopinto *et al*, 2004; Perez-Hernandez *et al*, 2013; Ramakrishnaiah *et al*, 2013; Sharma *et al*, 2011).

Because CD81 functions at the earliest stages of infection and at multiple points of cell-to-cell viral spread, targeting CD81 represents a distinct antiviral strategy. This approach complements DAAs and blocks host-dependent routes of viral transmission. While conventional biologics targeting mutable viral antigens remain vulnerable to viral escape, targeting a stable host entry factor like CD81 offers a higher genetic barrier to resistance. *De novo*-designed mini-proteins are well suited for this host-directed strategy. Their minimal size (∼40–80 amino acids), hyperstability, and customizable binding interfaces allow them to disrupt the protein–protein interactions required for viral entry (Cao *et al*, 2022; Huang *et al*, 2024; Ragotte *et al*, 2025b; Vazquez Torres *et al*, 2025). Designing a mini-protein that recognizes the CD81-LEL can sterically block E2–CD81 binding, disrupt receptor complex formation, and prevent both cell-free and cell-to-cell HCV transmission. CD81-targeting mini-proteins also address clinical scenarios where DAAs are insufficient. For example, they could serve as prophylactic agents to prevent graft reinfection following liver transplantation. In this setting, residual virus or infectious CD81-positive EVs rapidly infect the new allograft (Brown, 2005; Bush *et al*., 2014; Lin *et al*, 2015; Ramakrishnaiah *et al*., 2013). They also provide prophylaxis for high-risk individuals, such as healthcare workers or populations with repeated exposure (Houldsworth *et al*, 2014; Ji *et al*, 2015; Krieger *et al*, 2010; Meuleman *et al*, 2008). Given that CD81 facilitates the entry or budding of multiple pathogens—including HIV, diverse viral pathogens, and several *Plasmodium* species (e.g., *P. falciparum* sporozoite entry into hepatocytes)—CD81-targeting mini-proteins may offer broad-spectrum antiviral and antiparasitic utility (Al Olaby *et al*, 2014; Feneant *et al*, 2014; Silvie *et al*, 2003).

In this study, we describe the computational design and functional characterization of mini-19, a *de novo*-designed CD81-targeting mini-protein that inhibits HCV infection in cellular models. It was engineered to act as both a steric blockade against HCV attachment and a structural clamp. Supported by molecular dynamics (MD) simulations, we show that mini-19 restricts the fusogenic conformational plasticity of the receptor. This restriction presents a distinct mechanism of viral inhibition. These findings establish CD81-targeting mini-proteins as a class of host-directed entry inhibitors. By targeting an early step in the HCV life cycle, they offer broad potential as prophylactic and therapeutic agents that address the limitations of existing antiviral regimens.

## Methods

### *De novo* computational design of CD81-targeting mini-proteins

To engineer high-affinity viral entry inhibitors, we established a *de novo* mini-protein design pipeline targeting the CD81-LEL. The LEL coordinates from the full-length cryo-EM structure of CD81 (PDB: 5TCX) (Zimmerman *et al*, 2016) served as the computational template, with a focus on the C-and D-helices that constitute the binding platform for the HCV E2 glycoprotein. Using RFdiffusion (Watson *et al*, 2023), we generated *de novo* mini-protein backbones (60–80 amino acids) by conditioning the diffusion process to dock against a predefined hotspot interface (residues L162, V169, I182, and F186). To ensure structural stability and efficient folding, a three-helix bundle topology was specified. We modulated diffusion noise to control helical geometry during generation, setting both translational and rotational noise to 0.5. This balance allowed sufficient conformational exploration while maintaining stability. From an initial pool of 5,000 models, we selected 44 scaffolds based on RFdiffusion confidence metrics. Specifically, we filtered for a predicted Alignment Error (pAE) score < 5.99, indicating high structural certainty at the binding interface. Next, we optimized the sequences of these scaffolds using ProteinMPNN (Dauparas *et al*, 2022). For each scaffold, we performed 1,000 sequence design iterations to generate a diverse library of variants energetically compatible with the *de novo* backbones and the CD81 binding interface. We filtered leads based on a pAE score < 4.7 and validated the resulting models *in silico* using AlphaFold2 (AF2). A challenge in targeting the CD81-LEL arose from using a truncated soluble extracellular domain as the design template. This truncation exposed hydrophobic surfaces naturally shielded by the transmembrane domain in the full-length receptor. Consequently, several generated models exhibited non-specific binding to these exposed surfaces rather than to the target epitope. These mis-targeted candidates typically failed to converge, yielding low complex interface predicted Template Modeling (ipTM) scores ranging from 0.2 to 0.3. To eliminate these false positives, we applied a stringent multi-model selection threshold: a candidate was advanced for further validation only if at least two of the five independent AF2 models achieved an ipTM score > 0.83 while adopting identical backbone conformations. This structural convergence confirmed a single, well-defined binding mode at the intended CD81-LEL interface. Finally, to ensure independent folding stability, we required a predicted Local Distance Difference Test (pLDDT) score > 85 across the entire mini-protein sequence. Molecular graphics and structural analyses were performed using UCSF ChimeraX (Pettersen *et al*, 2021).

### Mini-protein expression and purification

Synthetic, codon-optimized genes encoding the mini-proteins (Gene Universal) were cloned into the pET-21a(+) vector, incorporating an N-terminal 8×His-tag and a TEV cleavage site. Proteins were expressed in *E. coli* BL21(DE3) by induction with 0.5 mM IPTG at an OD_600_ of ∼0.8, followed by overnight incubation at 18°C. Cells were lysed by sonication in Buffer A (20 mM Tris-HCl, pH 8.0, 300 mM NaCl, 5% glycerol) containing protease inhibitors. Clarified lysates were purified via TALON metal affinity chromatography (Takara Bio) and eluted with 300 mM imidazole in Buffer A. Eluates were either concentrated directly or subjected to TEV cleavage followed by passage through a Ni-NTA column (Qiagen) to remove the cleaved tag and TEV protease. Samples were further polished by size-exclusion chromatography using a Superdex 200 16/600 column equilibrated in 20 mM Tris-HCl (pH 8.0) and 300 mM NaCl. Monomeric peak fractions were pooled, and high purity was confirmed by SDS-PAGE. Final purified proteins were flash-frozen with 20% (v/v) glycerol and stored at −80°C. For biophysical characterization, His-tagged proteins were buffer-exchanged into 25 mM HEPES (pH 7.4) and 150 mM NaCl, or phosphate-buffered saline (PBS), using a PD MidiTrap G-25 column (Cytiva). All other downstream applications utilized samples formulated in 20 mM HEPES (pH 7.4) and 150 mM NaCl.

### Surface plasmon resonance (SPR) analysis

We characterized the binding affinities of the selected mini-proteins by SPR using a Biacore T200 system (Cytiva). All experiments were performed at 25°C using HBS-EP+ (10 mM HEPES, 150 mM NaCl, 3 mM EDTA, 0.05% Surfactant P20, pH 7.4) as the running buffer. The target protein, CD81-LEL (residues 113–201; Sino Biological, 14244-H07H), was immobilized onto the surface of a Series S sensor chip CM5 (Cytiva) via standard amine coupling. After pH scouting (pH 4.0 to 5.5), we selected 10 mM sodium acetate at pH 4.0 for immobilization. The target immobilization level was set to approximately 2,000 Resonance Units (RU) to ensure a sufficient maximum binding response (*R*_max_) for kinetic resolution. Binding interactions were monitored at a constant flow rate of 30 μL/min. We conducted an initial qualitative screening of the candidates by injecting each mini-protein at a fixed concentration of 1 μM. The association and dissociation phases were monitored for 60 s and 120 s, respectively. Candidates with significant binding responses were advanced to kinetic characterization using a 10-point, 2-fold serial dilution series starting from 1 μM. The dissociation phase was extended to 500 s to accurately resolve slow off-rates and determine the equilibrium dissociation constant (*K*_D_). Sensorgrams were recorded using Biacore T200 Control Software and processed with Biacore T200 Evaluation Software. We applied double-referencing to all data to account for non-specific binding and baseline drift. Finally, *K*_D_ values were calculated by fitting the processed data to a global 1:1 Langmuir binding model.

### Circular dichroism (CD) spectroscopy

We evaluated the secondary structure and conformational stability of the designed CD81 mini-proteins using far-UV CD spectroscopy on a J-1500 spectrometer (Jasco). Protein samples were prepared at 0.1 mg/mL in PBS. Wavelength scans were acquired from 190 to 260 nm at 25°C using a 1-mm path length quartz cuvette. Data collection parameters included a scanning speed of 50 nm/min, a digital integration time of 2 s, and a data pitch of 0.5 nm. Each spectrum represents the average of three technical accumulations with the corresponding buffer background subtracted. To assess thermal stability, we sequentially acquired CD spectra as the temperature increased from 25°C to 95°C at 10°C intervals (heating ramp: 7°C/min), maintaining identical scanning parameters. Raw ellipticity data were converted to mean residue ellipticity (MRE, [θ]) expressed in units of deg•cm^2^•dmol^-1^•res^-1^, for quantitative analysis.

### Molecular dynamics (MD) simulations

We performed atomistic MD simulations for the apo CD81-LEL and the CD81-LEL/mini-19 complex using GROMACS 2024.5 with the AMBER99SB-ILDN force field. To mimic the acidic endosomal environment crucial for viral entry, the protonation states of all titratable residues were assigned according to pH 4.0 prior to solvation. Each system was centered in a cubic box and solvated using the TIP3P water model, maintaining a minimum distance of 1.0 nm between the protein surface and the box edges. We neutralized the systems and added 150 mM NaCl to replicate physiological ionic strength. After energy minimization using the steepest descent algorithm, the systems were equilibrated in the NVT and NPT ensembles for 100 ps each, with positional restraints applied to the protein heavy atoms. During the production runs, the system temperature was maintained at 310 K using the stochastic dynamics integrator (Langevin dynamics) with an inverse friction constant (τ) of 2.0 ps. Pressure was controlled at 1 atm using the Parrinello-Rahman barostat with an isotropic coupling scheme. Following equilibration, we carried out unrestricted production simulations for 500 ns with a 2 fs time step. Periodic boundary conditions (PBC) were applied in all directions. Long-range electrostatic interactions were calculated using the particle mesh Ewald method with a 1.0 nm cutoff, and van der Waals interactions were truncated at the same distance. All covalent bonds involving hydrogen atoms were constrained by the LINCS algorithm. Each system was simulated in three independent runs with different initial velocities, utilizing GPU acceleration on an NVIDIA GeForce RTX 3090. Following the production runs, trajectory analysis was performed using the built-in analytical tools of the GROMACS package. Prior to analysis, all trajectories were structurally aligned to their initial conformations to remove PBC artifacts and global translational and rotational motions. We calculated the root mean square deviation (RMSD) and root mean square fluctuation (RMSF) of the protein backbone atoms to assess overall structural stability and local residue flexibility, respectively, across the three independent replicas.

### Flow cytometric binding assay using human PBMCs

Human peripheral blood mononuclear cells (PBMCs) were isolated from leukocyte reduction system chambers. For binding assays, 5 × 10⁵ PBMCs were first subjected to Fc receptor blockade for 20 min at 4°C to reduce non-specific binding. Cells were then incubated with His-tagged mini-proteins at 36°C for 20 min. Initial comparisons were conducted at 10, 100, and 1,000 nM, while the lead candidates, mini-19 and mini-20, were subjected to fine titration at concentrations ranging from 0.02 to 10 nM under the same conditions. After thorough washing with cold FACS buffer (PBS containing 2% FBS) to remove unbound ligands, cells were stained on ice with fluorochrome-conjugated antibodies against human CD19 (FITC, clone HIB19), His-tag (APC, clone J095G46), and human CD81 (PE, clone TAPA-1) (all from BioLegend). Following a final wash, samples were resuspended in FACS buffer and analyzed using a CytoFLEX flow cytometer (Beckman Coulter). Target engagement was quantified by evaluating the percentage of His-tag-positive events within the gated CD19⁺ B cell population.

### Preparation of fluorescent and biotinylated mini-proteins

To enable tracking in downstream cellular and EV binding assays, the N-termini of the mini-proteins were functionalized. Briefly, 60 nmol of each mini-protein in PBS (pH 7.0) was incubated with 1 mM Alexa Fluor 488 NHS ester or EZ-Link NHS-LC-Biotin (Thermo Fisher Scientific) for 12 h at 4°C with gentle agitation. The labeling reactions were quenched by adding 5 mM ethanolamine for 30 min. Unreacted reagents were subsequently removed using Zeba spin desalting columns (Thermo Fisher Scientific) pre-equilibrated with PBS (pH 7.4).

### HEK293 cell culture and generation of stable cell lines

Wild-type HEK293 (HEK293 WT, ATCC) and human CD81 knockout HEK293 (HEK293 hCD81 KO, Ubigene) cell lines were maintained in DMEM supplemented with 10% heat-inactivated FBS and 1% penicillin-streptomycin at 37°C in a 5% CO₂ humidified atmosphere. To generate cell lines for EV isolation, WT and KO cells were transduced with lentiviral vectors encoding a human CD81-GFP fusion protein (hCD81-GFP). Cells exhibiting the top 10% of GFP fluorescence intensity were flow-sorted to establish stable lines (HEK293 WT-hCD81-GFP and HEK293 hCD81 KO-hCD81-GFP).

### HEK293 cellular binding assay

To validate target specificity on living cells, HEK293 WT and HEK293 hCD81 KO cells were harvested and washed with serum-free DMEM. Approximately 1 × 10⁶ cells per reaction were incubated with Alexa Fluor 488-conjugated mini-proteins for 15 min at room temperature. Following incubation, cells were washed thoroughly with FACS buffer (PBS containing 1% FBS and 0.05% sodium azide), pelleted by centrifugation (300 × g, 5 min), and resuspended in FACS buffer. Target engagement was quantified by measuring cellular fluorescence using a CytoFLEX flow cytometer (Beckman Coulter).

### Isolation and purification of extracellular vesicles (EVs)

EVs were isolated following a previously established protocol (Choi *et al*, 2020). Briefly, the four established HEK293 cell lines were cultured to 80–90% confluency, washed with PBS and cultured in serum-free DMEM for 24 h. Conditioned media were harvested and cleared of cellular debris via sequential centrifugation (500 × *g* for 10 min, then twice at 2,000 × *g* for 15 min). The cleared supernatant was concentrated using a 100-kDa molecular weight cutoff hollow fiber membrane (GE Healthcare). For primary purification, the concentrate was layered onto 0.8 M and 2.0 M sucrose cushions and ultracentrifuged (SW32Ti rotor, Beckman Coulter) at 100,000 × *g* for 2 h at 4°C. The EV-containing interface was collected, diluted in HEPES-buffered saline (HBS; 20 mM HEPES, 150 mM NaCl), and adjusted to a final iodixanol concentration of 30%. This suspension was layered beneath a discontinuous iodixanol gradient (20% and 5%) and subjected to buoyant density ultracentrifugation (SW41Ti rotor, 200,000 × *g*, 2 h, 4°C). One-milliliter fractions were collected from the top, and the EV-enriched fractions were pooled. Total protein yield was determined by the Bradford assay (Bio-Rad Laboratories). Purified EVs were flash-frozen and maintained at −80°C for downstream applications.

### EV target engagement assay

To evaluate mini-protein target engagement on intact EVs, we utilized a customized chemiluminescence-based capture assay (Kim *et al*, 2017). High-binding 96-well plates were coated overnight at room temperature with an in-house rabbit polyclonal antibody targeting HEK293 whole-cell lysates. After blocking with 2% BSA in HBS, purified EVs (500 ng/mL) from the defined cell lines (HEK293 WT, WT-hCD81-GFP, hCD81 KO, and KO-hCD81-GFP) were captured on the plates. The captured EVs were then probed with biotinylated mini-19, followed by detection using streptavidin-horseradish peroxidase (SA-HRP). Extensive washing with PBST (PBS supplemented with 0.1% Tween-20) was performed between all steps. Upon the addition of a chemiluminescent substrate, luminescent signals were quantified using an EnSpire Alpha multimode plate reader (PerkinElmer).

### Super-resolution imaging of mini-19-bound EVs

Mini-19-bound EVs were visualized using the EV Profiler Kit (ONI) combined with direct stochastic optical reconstruction microscopy (dSTORM), with minor modifications. EVs were derived from placental stem cells (8.6 × 10⁹ particles/mL) and incubated with recombinant mini-19 protein (final concentration, 10 μM) for 2 h at 4°C to allow binding. EVs derived from placenta stem cells were used as a positive control without the binding step. For imaging, EV samples were affinity-captured onto microfluidic chips using antibodies against CD9, CD63, and CD81, which are canonical tetraspanin markers of EVs. To visualize mini-19, a His-tag–specific antibody conjugated to Alexa Fluor 647 was used in place of the CD81 antibody. Following capture, the dSTORM imaging buffer was freshly prepared and applied immediately before image acquisition. Super-resolution imaging was performed using a NanoImager S Mark II microscope (ONI) equipped with a 100× oil-immersion objective. Fluorescent signals corresponding to EV markers and mini-19 were acquired using 488 nm, 561 nm, and 647 nm laser lines, respectively. Data were reconstructed and analyzed using NimOS software (version 1.19.7, ONI) and further quantified using the ONI online analysis platform CODI (https://alto.codi.bio).

### Hepatoma cell lines and culture for HCV experiments

Huh-7.5 cells (Blight *et al*, 2002) and a Huh-7 derivative cell line (Koutsoudakis *et al*, 2007) were kindly provided by Dr. Charles M. Rice (Rockefeller University, New York, USA) and Dr. Ralf Bartenschlager (University of Heidelberg, Germany), respectively. Huh-7.5 cells were cultured and maintained in complete medium consisting of Dulbecco’s Modified Eagle Medium (DMEM; Welgene) supplemented with 10% fetal bovine serum (FBS; Gibco, USA), 1% non-essential amino acids (NEAA; Gibco, USA), 2 mM L-glutamine (Gibco, USA) and 100 units/mL penicillin and 100 µg/mL streptomycin (Gibco, USA). The Huh-7 derivative cell line was maintained in the same complete medium supplemented with 0.6 mg/mL Geneticin (G418; Gibco, USA).

### HCVcc production

Infectious cell culture-derived recombinant HCV particles (HCVcc) were produced as previously described (Lee *et al*., 2017). Briefly, the JFH1 E2p7-5A/5B-GFP reporter HCV construct was used to generate genomic HCV RNA transcripts. A total of 20 μg of HCV RNA transcripts was mixed with Huh-7 derivative cells (6 × 10^6^ cells) in a 0.4 cm gap cuvette and electroporated at 270 V and 975 μF using a GenePulser II (Bio-Rad, USA). HCVcc-GFP particles expressing an NS5A-GFP fusion protein were harvested at 2, 3, and 4 days post-transfection. The viral supernatant was clarified by filtration through a 0.22 μm syringe filter (Millipore, USA), concentrated using an Amicon Ultra centrifugal filter with a 100 kDa molecular weight cutoff (Millipore, USA), and stored at −80°C as viral stocks.

### HCVcc infection and assessment of antiviral activity

The antiviral activity of mini-proteins was assessed using an HCVcc-GFP infection system as previously described (Lee *et al*., 2017). Briefly, Huh-7.5 cells were seeded in 384-well plates at a density of 2.5 × 10^3^ cells per well. The following day, mini-proteins and reference inhibitors were serially diluted in complete DMEM and added to the cells 2 h prior to HCVcc inoculation. Cells were then infected with HCVcc-GFP and incubated at 37°C in a 5% CO_2_ incubator for 3 days. At 3 days post-infection (p.i.), the cells were fixed with 1% paraformaldehyde containing 10 μg/mL Hoechst 33342 (Life Technologies) for 1 h at room temperature. HCV infection was analyzed by determining the number of GFP-positive cells using a confocal microscope (Opera Phenix; PerkinElmer, USA). Cell viability was used as a marker for cytotoxicity by measuring the reduction in the cell population after drug treatment compared with untreated cells. Percent inhibition was calculated using the maximum infectivity of untreated cells as 0% inhibition and the background signal from infected cells treated with 10 µM sofosbuvir, a known HCV replication inhibitor, as 100% inhibition. The antiviral activity of mini-proteins and reference inhibitors was determined by dose-response curve (DRC) analysis and represented as the 50% inhibitory concentration (IC_50_) and the 50% cytotoxic concentration (CC_50_). All experimental conditions were tested in triplicate.

### Time-of-addition assay

To determine the stage of HCV infection affected by the compounds, a time-of-addition assay was performed as previously described (Lee *et al*., 2017) with minor modifications. Huh-7.5 cells were seeded in 384-well plates and treated with compounds (reference inhibitors or mini-proteins) at different time points before and/or after HCVcc infection. Compounds were applied under three different conditions as follows: Condition #1 (Continuous treatment), in which the compound was added 2 h prior to HCV infection and maintained throughout the entire experiment until fixation at 3 days post-infection (dpi); Condition #2 (Entry-stage treatment), in which the compound was added 2 h prior to HCV infection and maintained until 4 h post-infection, after which the cells were washed to remove the compound; Condition #3 (Post-entry treatment), in which the compound was added at 4 h post-infection, after removal of the virus inoculum, and maintained until fixation at 3 dpi. The antiviral activity of each compound was determined by DRC analysis as described above. All conditions were tested in triplicate (n = 3), except for the control conditions (untreated and 100% inhibition control), which were tested in ten replicates (n = 10). Reference inhibitors included an αCD81 antibody (BD Bioscience, USA), bafilomycin A1 (Sigma-Aldrich Korea), and sofosbuvir (Selleck Chemical, USA).

## Statistical analysis

All statistical analyses, dose-response curve fitting, and data visualization were performed using GraphPad Prism (GraphPad Software, Inc., USA). The number of independent replicates (n) for each experiment is specified in the respective figure legends or corresponding Methods sections. Where applicable, data are presented as the mean ± standard deviation (SD).

## Results

### Computational design of *de novo* mini-proteins targeting the CD81 large extracellular loop (LEL)

Our computational strategy aimed to competitively inhibit HCV entry by establishing a robust steric mask against HCV attachment and stabilizing the closed conformation of the CD81-LEL. At physiological pH (7.4), this predominant state is maintained by a core H151–Y127 hydrogen bond and a C157–C175 disulfide bridge (Cunha *et al*, 2017). During endosomal acidification, however, the protonation of key pH-antenna residues (D139 and E188) triggers a long-range redistribution of the hydration shell (Risueno *et al*, 2025). Together with the protonation of H151, this solvent-mediated destabilization drives a global shift toward the open, fusogenic conformation (Cunha *et al*., 2017; Risueno *et al*., 2025). To counter this, we designed mini-proteins specifically to physically occlude the viral epitope while simultaneously locking the head subdomain, thereby preventing the conformational transition required for viral fusion. To achieve this, we generated and rigorously filtered a vast library of *de novo* mini-proteins using a deep learning-based pipeline (detailed in the Methods). By applying stringent *in silico* confidence thresholds to eliminate non-specific binding artifacts and ensure independent folding stability, we successfully narrowed the initial pool down to 20 top-tier lead candidates. These were prioritized for recombinant expression and empirical validation (Fig. EV1 and Table EV1).

### Biophysical characterization of mini-protein binding to CD81

We successfully expressed and purified 15 of the 20 designed candidates to biochemical homogeneity, with robust expression yields reaching tens of milligrams per liter of culture. Tricine SDS-PAGE analysis confirmed their high purity and strict concordance with their expected molecular weights (Schagger & von Jagow, 1987) (Fig. EV2). To evaluate the binding kinetics and affinity of the purified mini-proteins for the CD81-LEL, initial screening was performed using surface plasmon resonance (SPR). At a concentration of 1 μM, eight constructs exhibited significant binding responses to the recombinant CD81-LEL, allowing determination of steady-state dissociation constants (*K*_D_) (Table EV2). Based on these binding profiles, mini-19 emerged as the primary lead candidate. The resulting sensorgrams were globally fitted to a 1:1 Langmuir binding model. Kinetic analysis revealed a fast association rate (*k*a = 7.38 x 10^5^ M^-1^s^-1^) and a slow dissociation rate (*k*d = 3.78 x 10^-4^s^-1^), yielding a *K*_D_ of 0.5 nM (Fig. 1A). The analyte was titrated over a concentration range from 31.3 nM down to 0.97 nM, closely bracketing the calculated *K*_D_ and robustly supporting the fit accuracy, confirmed by a low *χ*^2^ value of 36.4 RU^2^. The observed *R*_max_ of 1,431 RU represented ∼72% of the theoretical maximum capacity (2,000 RU), confirming that the immobilized CD81 retained its functional conformation without significant steric hindrance.

**Figure 1:**
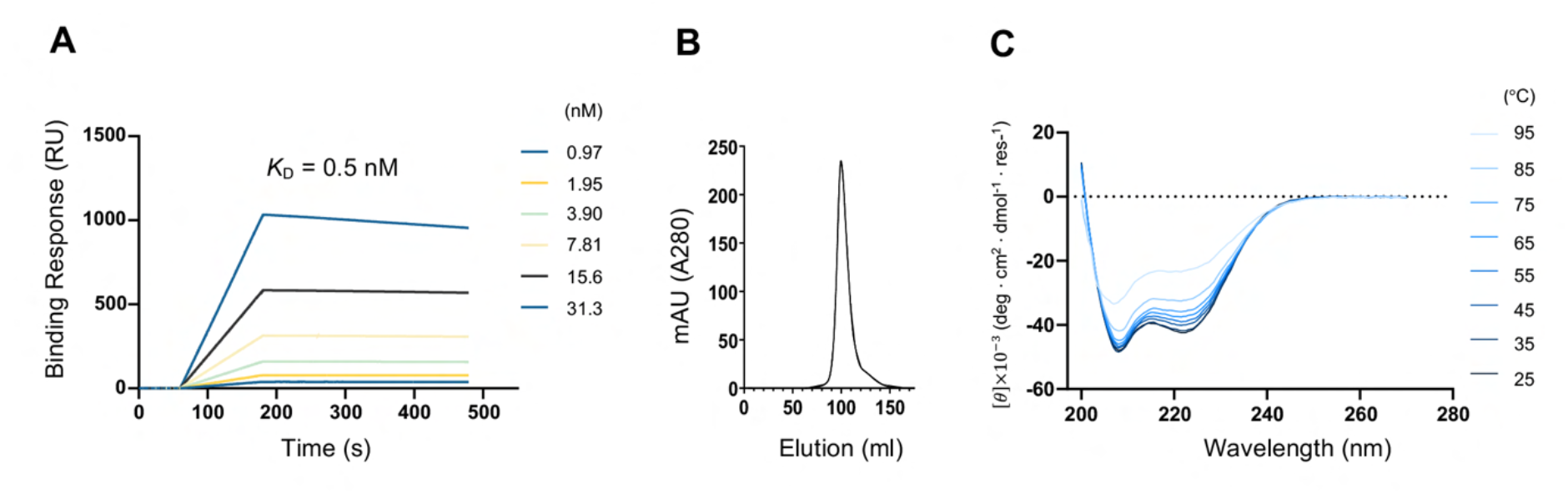
Biophysical characterization of the CD81 targeting mini-protein, mini-19. (**A**) Surface plasmon resonance (SPR) analysis demonstrating the binding affinity of mini-19 to the CD81-LEL. Representative SPR sensorgrams illustrate the dose-dependent binding of mini-19 to recombinant human CD81-LEL (n = 2). A Kinetic fitting to a 1:1 binding model was performed using data from analyte concentrations ranging from 0.97 nM to 31.3 nM, yielding a *K*D of 0.5 nM. (B) Size-exclusion chromatography profile of mini-19 on a Superdex 200 column, showing a single elution peak indicative of a monodisperse monomeric state. (C) Representative far-UV circular dichroism (CD) spectra of mini-19. The y-axis represents mean residue ellipticity ([θ]). The presence of two negative bands at 208 nm and 222 nm confirms an alpha-helical fold. This alpha-helical signature is retained at temperatures up to 95°C.

Interestingly, mini-20, which shares the same scaffold, exhibited attenuated affinity (*K*_D_ of 24 nM), highlighting the impact of subtle sequence variations on interface stability. Size-exclusion chromatography confirmed that mini-19 exists as a stable, homogeneous monomer in solution (Fig. 1B and Fig. EV3). Circular dichroism (CD) spectroscopy further confirmed that these lead candidates are highly thermostable and retain their designed α-helical secondary structure, consistent with the *de novo* design (Fig. 1C). Following the guidelines established by Greenfield (Greenfield, 2006), a reliable determination of the melting temperature requires a complete thermal transition profile, including a defined baseline for the fully unfolded state. Notably, consistent with the extreme thermal resilience characteristic of computationally designed mini-proteins (Cao *et al*., 2022; Huang *et al*., 2024; Ragotte *et al*., 2025b; Vazquez Torres *et al*., 2025), our scaffolds proved to be hyperstable; they retained their strong helical signatures at temperatures up to 95°C without exhibiting a cooperative unfolding transition. These results demonstrate that the *de novo-*designed mini-proteins possess the exceptional biophysical stability required for potential therapeutic applications under diverse physiological and environmental conditions.

### Structural basis of the CD81–mini-19 interaction interface

As a primary validation of our computational design, we evaluated the *in silico* confidence metrics of the generated candidates. The lead candidate, mini-19, exhibited a well-folded unbound state (monomer pLDDT = 94). Furthermore, the complex prediction yielded a multimer ipTM of 0.90 and a low pAE interaction score of 4.15, indicating robust target engagement. Subsequent *in silico* structural analysis suggested the basis for these metrics: mini-19 is predicted to adopt a topology that effectively covers the CD81-LEL target region, while the 20 selected candidates featured diverse structural scaffolds. For instance, the non-functional control mini-13 adopted a simple parallel three-helix bundle that failed to sufficiently bury the target epitope. In contrast, mini-19 features a more three-dimensional architecture where its first α-helix is stacked on top of the second and third helices. This topology allows mini-19 to more extensively wrap around and embrace the target region, effectively capping the binding interface. Surface representation colored by hydrophobicity reveals that this targeted region on the CD81-LEL constitutes a distinct hydrophobic patch (Fig. 2A). To further elucidate the spatial coordination of this interface, we utilized custom-coded SVG helical projections to illustrate the dual-function amphipathic core (Fig. 2B). Notably, the cross-sectional mapping demonstrates how the hydrophobic residues of Helix 1 serve a dual role: anchoring the structural core of the mini-protein while simultaneously participating in hydrophobic engagement with the receptor. By engaging the conserved helical bundle of the CD81-LEL, this structural arrangement is predicted to facilitate an extensive hydrophobic network. Rather than a single contiguous patch, this interaction is modeled to be driven by strong van der Waals packing between the CD81-LEL (L162, L165, V169, I182, L185, and F186) and specific hydrophobic clusters distributed across all three helices of mini-19: Helix 1 (L13, V17, L20, V21), Helix 2 (L36, I39, I43, L46), and Helix 3 (L61, I64) (Fig. 2C). Importantly, this extensive hydrophobic network is strictly confined to the structural core and the binding interface; the solvent-exposed exterior of mini-19 is predominantly hydrophilic, consistent with the high solubility and lack of aggregation observed in our SEC analysis (Fig. 1B and Fig. EV3). Crucially, this interface is predicted to tightly engage F186, a key molecular determinant for HCV E2 glycoprotein binding (Drummer *et al*, 2002; Higginbottom *et al*., 2000; Meola *et al*, 2000). This hydrophobic core is further stabilized by two hydrogen bonds: a peripheral interaction between E42 of mini-19 and S160 of CD81-LEL, and a buried interaction between T57 of mini-19 and T166 of CD81-LEL.

**Figure 2:**
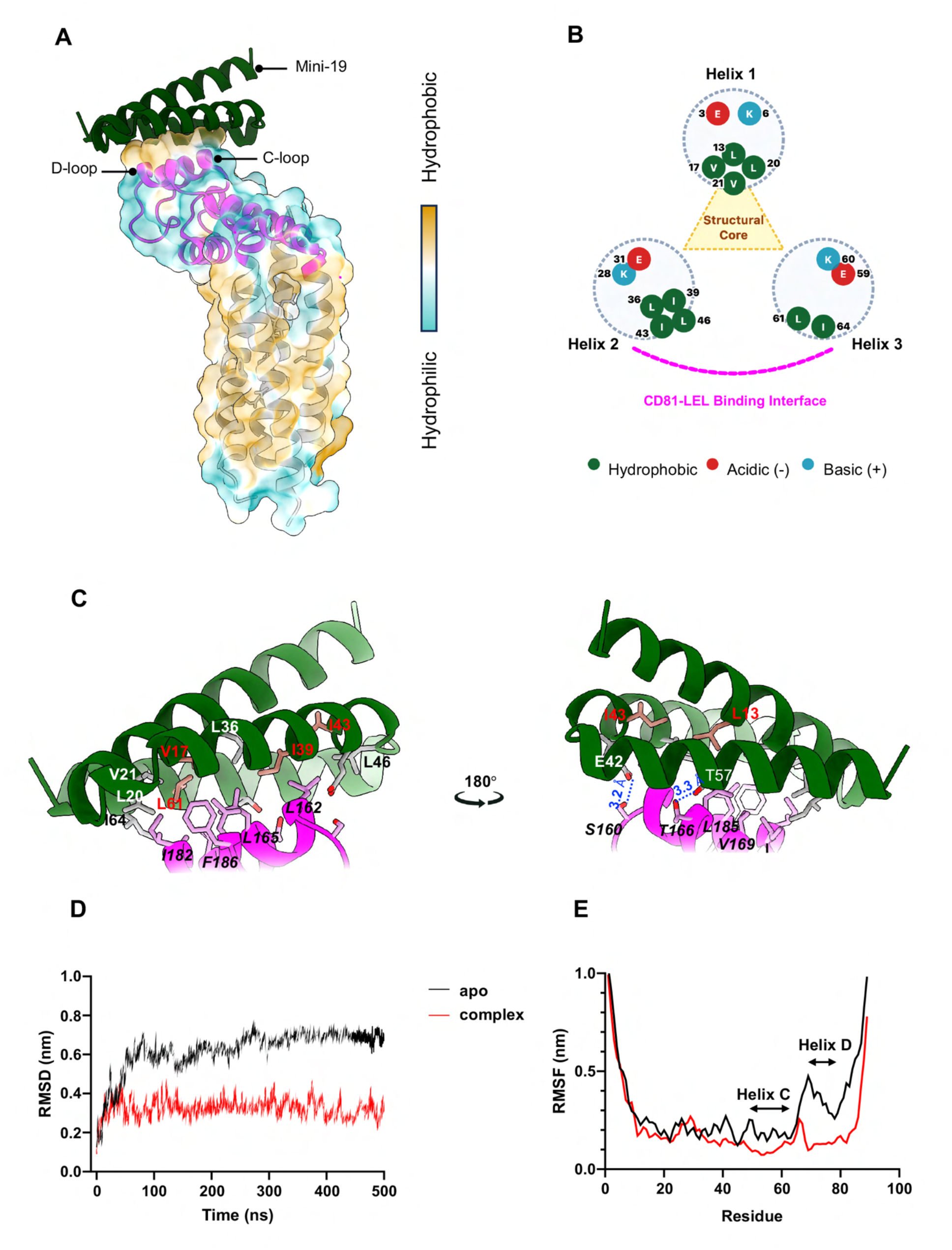
Predicted structural basis of the interaction between mini-19 and the CD81 receptor. (**A**) Overall AlphaFold2 (AF2)-predicted model of the *de novo* designed mini-19 (forest-green) in complex with CD81. CD81 is displayed as a hydrophobicity-colored, semi-transparent surface (50%), with the LEL domain highlighted in magenta. (**B**) Detailed cross-sectional schematic of the mini-19 binding interface. Helical projections, generated via custom-coded visualization scripts, illustrate the dual-function amphipathic core. Residues in Helix 1 form the structural core (yellow triangle) while simultaneously contributing to the extensive hydrophobic network at the CD81-LEL binding interface (magenta dashed line), in coordination with strategic hydrophobic clusters on helices 2 and 3. Residues are color-coded by physicochemical properties: dark green (hydrophobic), red (acidic), and cyan (basic). (**C**) Detailed atomic views of the predicted interaction interface. Mini-19 engages the LEL domain of CD81 predominantly through clustered hydrophobic and van der Waals interactions. Two key hydrogen bonds are modeled to stabilize the complex: one at the interface periphery between E42 (mini-19) and S160 (CD81-LEL), and another within the hydrophobic pocket between T57 (mini-19) and T166 (CD81-LEL). Amino acid residues involved in the interaction interface are indicated in italics for CD81-LEL. Residues selected for binding-deficient mutant generation are highlighted in brown with red characters. Molecular graphics in (A) and (C) were generated using UCSF ChimeraX. **(D)** Comparison of the backbone RMSD trajectories over a 500 ns MD simulation, performed at an acidic endosomal pH (pH 4.0), for the apo state (black line) and the mini-19 complex state (red line). The plotted trajectories represent one representative simulation selected from three independent replicas. The complex state maintains lower and more stable RMSD values (∼0.3–0.4 nm) compared to the unbound apo state (∼0.6–0.7 nm), indicating that mini-19 restricts the overall structural fluctuation of CD81-LEL. **(E)** Comparison of the RMSF profiles between the apo state (black line) and the mini-19 complex state (red line) during the MD simulations at pH 4.0. The plotted profiles represent data derived from the same representative simulation selected from three independent replicas. The binding of mini-19 restricts the overall structural flexibility of CD81-LEL, with stabilization observed in the dynamic regions spanning helices C and D.

To evaluate the structural stability of the modeled interface under the fusogenic endosomal environment, we analyzed the CD81-LEL/mini-19 complex using 500 ns MD simulations at pH 4.0. Mini-19 binding significantly reduced the overall RMSD compared to the apo state (Fig. 2D, Fig. EV4). While the apo-LEL exhibited high flexibility under this acidic stress, the complex remained distinctly constrained. Furthermore, marked decreases in RMSF across helices C and D indicated suppressed conformational dynamics within the LEL region (Fig. 2E, Fig. EV5). These results demonstrate that mini-19 acts as a structural clamp, restricting the receptor’s plasticity even under fusion-permissive conditions.

### Experimental validation of the binding interface via targeted mutagenesis

To validate the modeled interface, we performed targeted combinatorial mutagenesis on five core contact residues (L13, V17, I39, I43, and L61). We generated two variants: the 5A mutant (L13A/V17A/I39A/I43A/L61A) and the 4A1E mutant (incorporating a V17E charge-reversal substitution). SPR analysis revealed a substantial loss of affinity for both mutants compared to the parental mini-19 (*K*_D_ = 0.5 nM). The 5A mutant exhibited severely reduced affinity (*K*_D_ = 0.88 mM), while the 4A1E mutant nearly abolished binding (*K*_D_ = 28.5 mM) (Fig. EV6). This multi-log decrease in affinity provides compelling functional evidence that these specific residues constitute the energetic hotspot of the complex, confirming the precision of our *in silico* model. To ensure that the loss of binding was strictly due to interface disruption rather than global unfolding, we assessed the conformational integrity of the mutants using size-exclusion chromatography. Purified mini-19 and its binding-deficient mutants were analyzed on a Superdex 75 Increase 10/300 GL column. None of the proteins showed misfolding-induced aggregation (Fig. EV3). The parental mini-19 eluted as a sharp monomeric peak consistent with its theoretical mass of 10.2 kDa. In contrast, mutants 5A and 4A1E eluted slightly earlier, corresponding to apparent masses of ∼11.3 kDa and ∼11.6 kDa, respectively. Although slightly elevated, these masses remained well resolved from the expected dimeric elution volume (∼12.2 mL). This shift indicates an increased hydrodynamic radius, suggesting minor conformational expansion likely due to decreased packing density within the hydrophobic core. Furthermore, far-UV CD spectroscopy confirmed that both variants retained a predominantly α-helical signature (Fig. EV7). A moderate shift in molar ellipticity at 222 nm likely reflects local helical reorientation to accommodate the modified core packing, rather than global destabilization. The preservation of the overall spectral profile across a broad temperature range (up to 95°C) demonstrates that the global fold remains intact. Together, these data confirm that the loss of CD81 engagement is directly attributable to the specific ablation of interface residues.

### Physiological CD81 engagement by mini-19

To evaluate the capacity of mini-19 to engage its target in a physiological context, we performed flow cytometric analysis on human PBMCs (Fig. 3A). Mini-19 demonstrated robust, specific binding to CD19⁺ B cells, a lymphocyte subpopulation well-characterized by high endogenous surface expression of CD81. This interaction strictly depended on the designed interface: while mini-19 and its structural ortholog (mini-20) strongly engaged the target, the flat-topology control (mini-13) showed no detectable binding (Fig. EV8).

**Figure 3:**
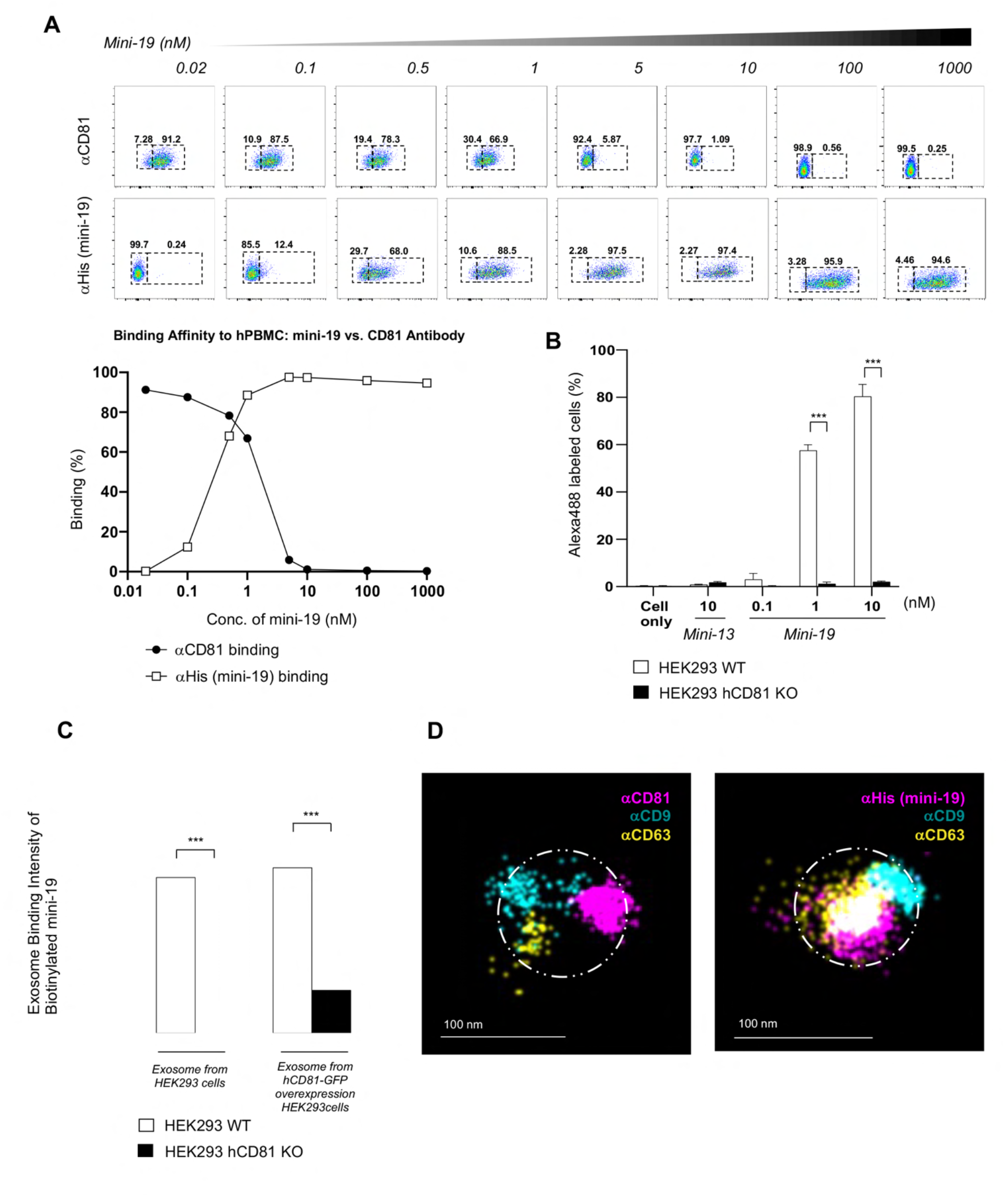
Mini-19 specifically engages CD81 in physiological contexts on human cells and extracellular vesicles (EVs). (**A**) Receptor occupancy and epitope blocking by mini-19 on primary human B cells. Primary human PBMCs were incubated with increasing concentrations of His-tagged mini-19 (0.02 nM to 1,000 nM), washed, and stained with a fluorescently labeled αCD81 antibody. Flow cytometric analysis of the CD19⁺ B cell population demonstrates a dose-dependent increase in mini-19 binding (His-tag detection, bottom row) that coincides with the progressive loss of the CD81 antibody signal (top row). This reciprocal signal shift confirms that pre-bound mini-19 occupies the CD81 receptor, masking the epitope and preventing subsequent antibody engagement. **(B)** Binding of mini-proteins to wild-type (WT) and hCD81-knockout (KO) HEK293 cell lines. Following incubation with Alexa Fluor 488-labeled mini-19 or the control mini-13, the percentage of fluorescent cells was quantified by flow cytometry. **(C)** Binding of mini-19 to EVs derived from four distinct HEK293 cell lines (WT, hCD81-GFP, hCD81 KO, and hCD81 KO-hCD81-GFP). EVs were assayed at equivalent concentrations (500 ng/mL, 0.1 mL/well). An in-house αHEK293 whole-cell lysate polyclonal antibody was used for capture, and biotinylated mini-19 was utilized for detection. For (B) and (C), data are presented as mean ± SD (n = 2). ***P < 0.001, calculated by two-way ANOVA with Bonferroni correction for multiple comparisons. (**D**) Super-resolution imaging of control and mini-19-bound EVs. Placental stem cell–derived EVs were captured using the EV Profiler Kit (ONI). EVs were affinity-captured on microfluidic chips using antibodies against the canonical tetraspanin markers CD9, CD63, and CD81 (left). Bound mini-19 was detected using an Alexa Fluor 647-conjugated αHis-tag antibody (right). Representative super-resolution images demonstrate the spatial colocalization of mini-19 with established EV markers. White dashed circles indicate the estimated EV size based on typical EV dimensions. Scale bar, 100 nm.

Pre-incubation of human PBMCs with escalating concentrations of mini-19 resulted in a dose-dependent reduction of subsequent staining by a commercial αCD81 antibody. This competitive inhibition demonstrates that mini-19 specifically binds the native CD81 receptor on living cells, sterically masking the critical epitope and preventing antibody engagement. To further validate target specificity, we performed binding assays using HEK293 cells, an established model for native CD81 expression, as a complementary cellular system. Mini-19 exhibited a clear dose-dependent increase in binding signal to HEK293 WT cells, whereas the negative control, mini-13, showed no detectable interaction. Critically, the binding signal was abolished in HEK293 hCD81 KO cells, providing definitive genetic evidence that the cellular uptake or surface attachment of mini-19 is mediated exclusively through the CD81 receptor (Fig. 3B). Beyond cell-surface engagement, we investigated the specificity of mini-19 for CD81 on EVs, which serve as key vehicles for receptor-assisted viral dissemination. Biotinylated mini-19 was incubated with EVs derived from both wild-type (WT) and CD81-knockout (KO) HEK293 cells. Strong binding signals were observed exclusively in WT-derived EVs, with a complete loss of signal in the KO model (Fig. 3C). This result confirms that mini-19 interacts specifically with the CD81 receptor and lacks cross-reactivity with other abundant EV surface proteins. To visualize this precise target engagement at the nanoscale, we employed direct stochastic optical reconstruction microscopy (dSTORM) on EVs derived from placental stem cells. Control EVs exhibited a canonical distribution of the tetraspanin markers CD9, CD63, and CD81 (Fig. 3D, left), whereas vesicles incubated with mini-19 displayed dense, specific localization of the mini-protein on their surface (Fig. 3D, right). Importantly, the mini-19 signal exhibited spatial colocalization with CD9 and CD63 within the ∼100 nm vesicular boundary. This high-density clustering demonstrates that mini-19 accurately targets the native CD81 receptor even when embedded within complex tetraspanin-enriched microdomains (TEMs). Collectively, these nanoscale imaging data—combined with our cellular binding assays—establish that mini-19 precisely recognizes the physiological fold of CD81 within highly complex native membranes.

### Potent HCV antiviral efficacy and entry-specific inhibition by mini-19

We evaluated the antiviral efficacy of six computationally designed mini-proteins (mini-3, 4, 7, 16, 19, and 20) against HCVcc infection in Huh-7.5 cells (Fig. EV9). Mini-19 exhibited the most potent antiviral activity, consistent with its superior binding affinity observed in our biophysical assays (Fig. 1A, Table EV2). Mini-20 demonstrated comparable, albeit slightly attenuated, binding and antiviral efficacy. In contrast, the remaining candidates (mini-3, 4, 7, and 16) showed negligible antiviral effects, and the structurally distinct negative control, mini-13, lacked antiviral activity. These results highlight a strong correlation between the binding affinity (*K*_D_ values) determined by SPR and cellular antiviral activity. Specifically, the lead candidate, mini-19, demonstrated potent, dose-dependent inhibition with an IC_50_ of 1.2 nM (Fig. 4), comparable to or superior to previously reported αCD81 monoclonal antibodies (Desombere *et al*, 2017; Fofana *et al*, 2013). To confirm that this antiviral effect was mediated through targeted antagonism of the CD81-LEL helical bundle, we evaluated the binding-deficient mutants 5A and 4A1E. Fluorescence microscopy and dose-response curve (DRC) analysis revealed a dramatic loss of inhibitory capacity for these mutants, marked by a > 3-log increase in IC_50_ values (approximately 4.5 µM for 5A and 3.5 µM for 4A1E) compared with the parental mini-19 (0.5 nM) (Fig. 4). Furthermore, all candidates maintained high cell viability (CC_50_ >10 µM), indicating that the observed antiviral effects were not driven by cytotoxicity (Figure. 4B, C). Collectively, these results establish that the potent antiviral efficacy of mini-19 is strictly dependent on its precise structural engagement with the CD81 receptor.

**Figure 4:**
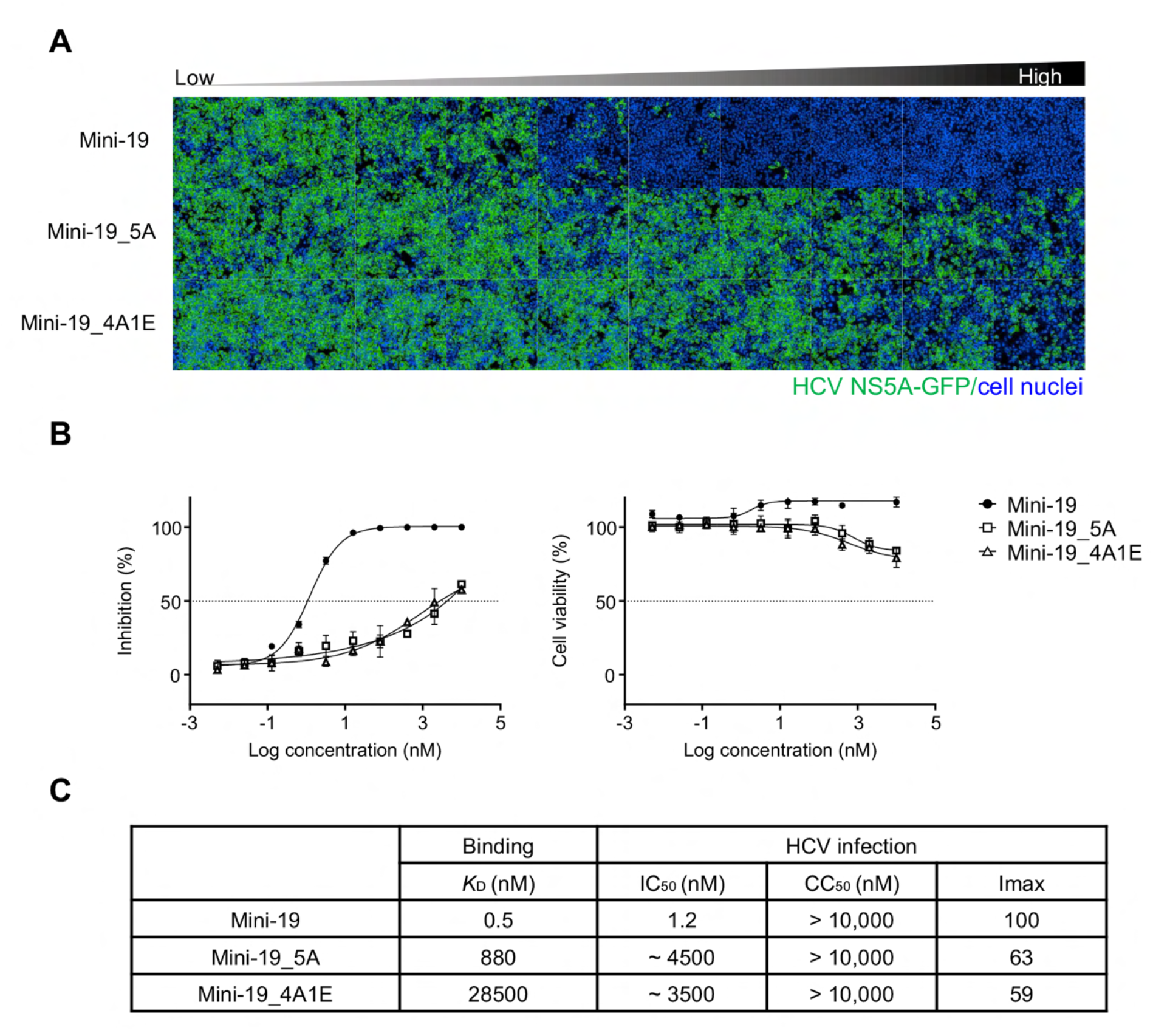
Antiviral efficacy of *de novo* designed CD81-targeting mini-19 and its mutants against HCVcc infection. Mini-19 and two binding-deficient mutants (5A and 4A1E), were tested at ten different concentrations, starting at a maximum concentration of 5 µM followed by 5-fold serial dilutions. Huh-7.5 cells were pre-treated with the compounds for 2 h prior to HCV infection. The cells were then inoculated with 40 µl of HCVcc-GFP (JFH1-E2p7-NS5A-GFP) and incubated for 72 h. At 3 days post-infection (dpi), the cells were fixed with 1% paraformaldehyde containing Hoechst 33342 (1:1000 dilution) to stain nuclei. HCV infection was visualized by measuring GFP signal using confocal microscopy. (**A**) Representative images from one of nine subfields acquired per well at each concentration are shown. All conditions were tested in triplicate. Green indicates HCV infection (GFP), and blue indicates cell nuclei (Hoechst). **(B)** Dose-response and cell viability curves. Percent inhibition was normalized relative to untreated infected cells (0% inhibition) and cells treated with 10 µM sofosbuvir (100% inhibition). Cell viability was quantified based on nuclei counts. Black circles, white squares, and white triangles indicate mini-19, 5A, and 4A1E, respectively. (**C**) Summary of antiviral activity. Half maximal inhibitory concentration (IC50) and Half maximal cytotoxic concentration (CC50) were determined by dose response curve (DRC) analysis. Imax is maximum inhibition.

### Elucidation of mini-19 as a competitive antagonist of initial viral attachment

To define the precise stage of the HCV life cycle targeted by mini-19, we performed time-of-addition assays using the HCVcc infection system, including both entry and replication inhibitors as references (Fig. 5). The replication inhibitor sofosbuvir achieved near-complete inhibition (Imax = 100 %) across all treatment regimens (Conditions #1, #2 and #3).

**Figure 5:**
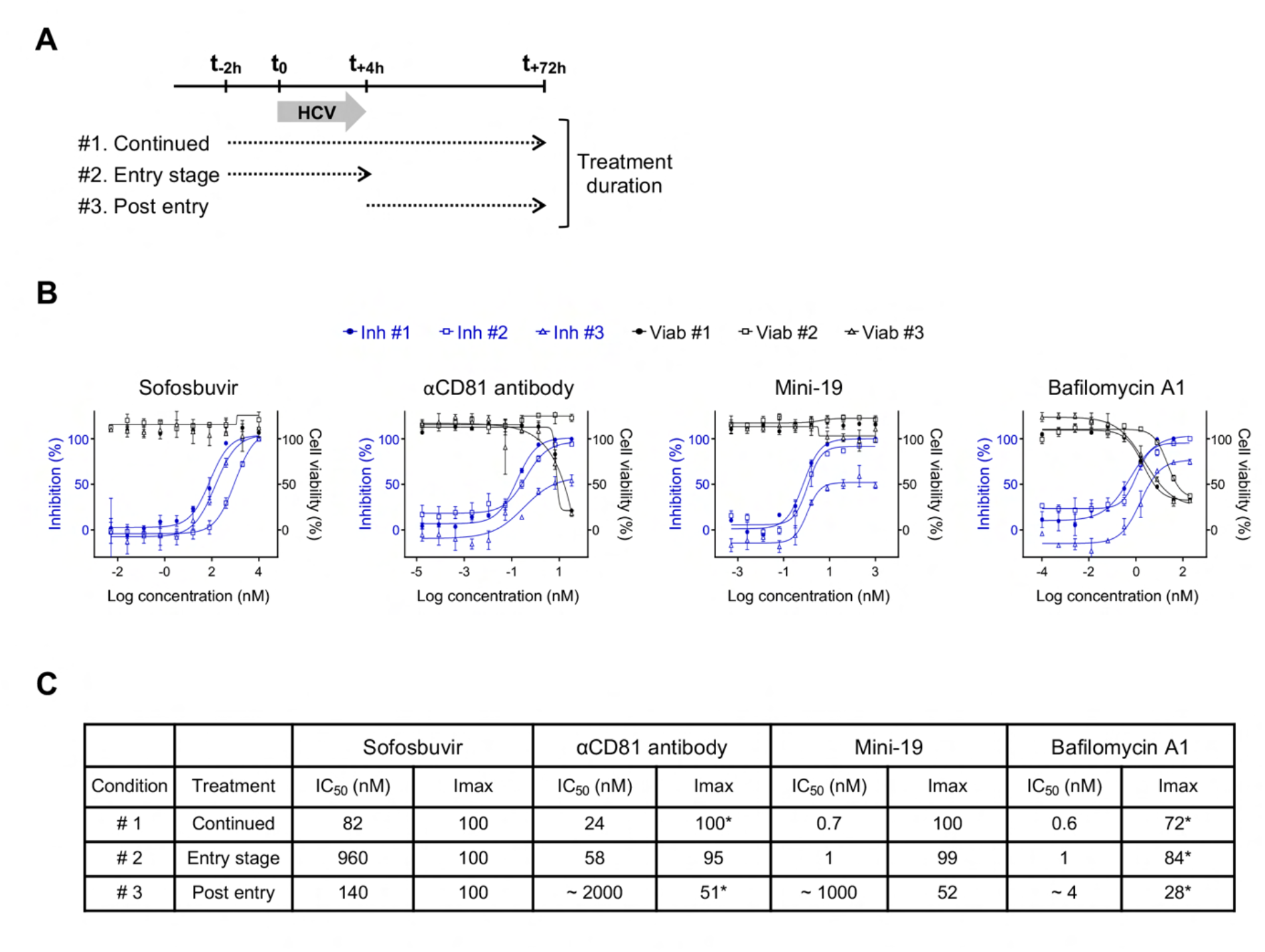
CD81-targeting mini-19 inhibits HCV infection at the viral entry stage. (A) Schematic diagram of the time-of-addition assay used to identify the inhibitory stage of mini-19 during HCVcc infection. Huh-7.5 cells were treated with compounds under three different regimens relative to HCVcc inoculation (t0) as indicated: (#1) continuous treatment, in which the compound was added at t-2h and maintained until t+72h; (#2) entry-stage treatment, in which the compound was added at t-2h and removed at t+4h; and (#3) post-entry treatment, in which the compound was added at t+4h after removal of input virus and maintained until t+72h. In all conditions, the viral inoculum was removed at t+4h, and HCV infection was determined by measuring GFP-positive cells using confocal microscopy at 72 hours post-infection. (**B**) Dose-response curve (DRC) analysis of mini-19 and reference inhibitors. The compounds were evaluated using 10-point, 5-fold serial dilutions. The highest concentrations were 33 nM for αCD81 antibody, 200 nM for bafilomycin A1, 10 µM for sofosbuvir, and 1 µM for mini-19. Quantitative data were normalized and plotted. Circles, squares and triangles indicate treatment Conditions #1, #2 and #3, respectively. Blue lines indicate percent inhibition, and black lines indicate percent cell viability. (**C**) Summary of IC50 and Imax values derived from the DRC analysis. *Imax values were determined only for conditions maintaining > 75% cell viability.

Notably, even under a limited 6-h pulse treatment (Condition #2, 2 h pre-and 4 h post-infection), sofosbuvir maintained its potency (IC_50_ = 82 nM), consistent with the efficient intracellular uptake and long half-life of its active metabolite (Murakami *et al*, 2010). In contrast, mini-19 exhibited a time-dependent inhibition pattern that closely paralleled that of the αCD81 antibody. Specifically, entry-stage treatments (Conditions #1 and #2) with mini-19 provided robust protection against HCV infection, yielding IC_50_ values of 0.7 nM and 1nM, respectively. However, delayed treatment (Condition #3) at 4 h post-infection, following the removal of the viral inoculum, resulted in a nearly 1,000-fold increase in the IC_50_ to approximately 1 µM and a significant reduction in Imax to 52% (Fig. 5C). This marked loss of potency post-viral entry indicates that mini-19 primarily targets the early, extracellular stages of HCV infection. To further isolate the mechanism of viral entry, we compared mini-19 with bafilomycin A1, a V-ATPase inhibitor that prevents endosomal acidification. While both the αCD81 antibody and mini-19 exhibited a dramatic 2-to 3-log shift in IC_50_ between Conditions #1 and #3, bafilomycin A1 showed a relatively modest shift (6.6-fold) between these conditions. Furthermore, during early treatment (Condition #2), both the αCD81 antibody and mini-19 achieved near-complete inhibition, whereas bafilomycin A1 displayed a reduced Imax (∼85%). The particularly low Imax observed for bafilomycin A1 in Condition #3 is attributed to the exclusion of high-concentration data points where cell viability dropped below 80% due to compound-induced cytotoxicity. Collectively, these data confirm that bafilomycin A1 acts at a later stage of entry than CD81-targeting agents. Crucially, while mini-19 shares a nearly identical stage-specific inhibitory profile with the αCD81 antibody—confirming its role as an antagonist of the early HCV attachment phase—it achieves this targeted blockade with strikingly enhanced potency, exhibiting an IC_50_ profile that vastly outperforms the monoclonal antibody. Finally, the antiviral activity of mini-19 is highly specific to the HCV-CD81 interaction, as it exhibited no significant activity against other flaviviruses, such as Zika virus (ZIKV) and Dengue virus (DENV), which utilize CD81-independent entry pathways (Fig. EV10).

## Discussion

The development of *de novo* mini-proteins targeting host factors marks a paradigm shift in antiviral strategy, transcending the limitations of traditional small-molecule inhibitors and monoclonal antibodies. Unlike conventional biologics targeting mutable viral antigens—which are inherently vulnerable to viral escape—host-directed therapies offer a substantially higher genetic barrier to resistance. In this study, we engineered mini-19, a sub-nanomolar mini-protein (*K*_D_ = 0.5 nM) that targets the CD81-LEL, demonstrating potent inhibition of HCV entry.

The structural rationale for targeting the CD81-LEL is rooted in its role as a central organizer of the viral entry complex. Computational design specifically targeted the C- and D-helices of the LEL, a region of significant conformational plasticity. Mutagenesis and structural studies have long identified F186 and I182 as indispensable determinants for the interaction with the viral E2 glycoprotein (Drummer *et al*., 2002; Higginbottom *et al*., 2000; Meola *et al*., 2000; Rajesh *et al*, 2012). By docking mini-19 into this helical pocket, we hypothesized and subsequently confirmed the steric occlusion of the E2-binding site. The multi-log loss of affinity observed in our 5A and 4A1E mutants (*K*_D_ of 880 nM and 28.5 μM, respectively) confirms that the potency of mini-19 is derived from precise hydrophobic complementarity at this conserved hotspot. Furthermore, our strategy of targeting the closed conformation of CD81—stabilized by a core H151-Y127 hydrogen bond and a C157-C175 disulfide bridge (Cunha *et al*., 2017; Risueno *et al*., 2025)—provides a mechanistically distinct approach. By locking the receptor in its non-fusogenic state, mini-19 pre-emptively blocks the conformational transition required for membrane fusion in the acidic endosomal environment.

While monoclonal antibodies targeting CD81 show prophylactic potential, their clinical utility is constrained by their large molecular size (∼150 kDa), high production costs, and limited tissue penetration. At ∼8 kDa, mini-19 is nearly 20-fold smaller than standard monoclonal antibodies, offering significant advantages in tissue penetration and molar density at the infection site. This is particularly relevant in the context of liver transplantation, where rapid penetration into the hepatic parenchyma is essential to prevent graft reinfection from residual virus or CD81-positive EVs (Brown, 2005; Bush *et al*., 2014; Lin *et al*., 2015; Ramakrishnaiah *et al*., 2013). Additionally, the high thermostability of mini-19 (up to 95°C) eliminates the need for cold-chain logistics, a distinct advantage for global health applications. Beyond direct viral inhibition, the specific engagement of mini-19 with EV-associated CD81 opens broader therapeutic avenues. We demonstrated that mini-19 recognizes the native topology of CD81 on EVs, co-localizing with the canonical EV markers CD9 and CD63 in dSTORM super-resolution imaging. Given that tumor-derived EVs exploit CD81-mediated uptake to promote epithelial–mesenchymal transition and prepare the pre-metastatic niche (Bailly & Thuru, 2023; Zhou *et al*, 2021), mini-19 could serve as a biologic shield against oncogenic EV crosstalk (Susa *et al*, 2020). Furthermore, mini-proteins offer the versatility of intracellular delivery or *in situ* expression, enabling sustained receptor blockade without repeated systemic dosing.

Despite the promising *in vitro* profile, the successful deployment of this mini-protein *in vivo* requires addressing pharmacokinetic limitations. Because its size (∼8 kDa) is below the glomerular filtration threshold (∼60 kDa), mini-19 is susceptible to rapid renal clearance and possesses a limited serum half-life. Thus, the present study serves as a foundational validation of the binding interface. To circumvent rapid clearance, future iterations must incorporate half-life extension modalities, such as Fc-fusion or albumin-binding modules. Notably, the rigid, highly packed hydrophobic core of our computationally designed scaffold is expected to confer superior resistance to proteolytic degradation compared with flexible peptides. Building upon pioneering advancements in *de novo* protein design, such hyperstable scaffolds have been further engineered to withstand extreme physiological conditions, including gastric acid and severe proteolytic environments, facilitating potential oral administration (Berger *et al*, 2024; Cao *et al*, 2020; Ragotte *et al*, 2025a).

A primary concern in targeting a ubiquitous host factor is the potential for systemic toxicity; however, multiple lines of evidence suggest a favorable safety profile for transient CD81 blockade. Although CD81 participates in B-cell activation (Bailly & Thuru, 2023; Susa *et al*, 2021; Susa *et al*., 2020), hepatocyte membrane organization (Feneant *et al*., 2014; Reynolds *et al*, 2008), and tetraspanin-mediated signaling (Brazzoli *et al*., 2008; Carloni *et al*, 2004; Cherukuri *et al*, 2004; Gerold *et al*., 2020; Hammerstad *et al*, 2017; Pan *et al*, 2024), the expected side effects of inhibition appear minimal. CD81-deficient mice develop normally (Maecker & Levy, 1997; Miyazaki *et al*, 1997) and exhibit only mild alterations in B-cell receptor signaling (Tsitsikov *et al*, 1997), suggesting that short-term inhibition is unlikely to cause significant immunological dysfunction (Bailly & Thuru, 2023; Vences-Catalan *et al*, 2019). Hepatocyte-specific roles of CD81 also appear dispensable for liver architecture and function (Ding *et al*, 2017; Miyazaki *et al*., 1997; Zimmerman *et al*., 2016). Importantly, CD81 inhibition may even reduce the entry efficiency of multiple pathogens that exploit TEMs for host cell invasion (Florin & Lang, 2018; Martin *et al*, 2005; Monk & Partridge, 2012), including HIV and *Plasmodium* species (Al Olaby *et al*., 2014; Feneant *et al*., 2014; Silvie *et al*., 2003; Zona *et al*, 2014). In the context of HCV, targeting the CD81-LEL is a therapeutically advantageous strategy for blocking both cell-free and cell-to-cell viral propagation (Ahmed *et al*, 2021; Aljowaie *et al*, 2020; Bailly & Thuru, 2023; Fofana *et al*., 2013; Rajesh *et al*., 2012; Zhang *et al*, 2004; Zona *et al*., 2014).

While our data highlight the potent neutralizing capacity of mini-19, predicting its exact *in vivo* efficacy remains challenging. Standard HCVcc models using immortalized hepatoma cell lines may not fully recapitulate the repertoire and spatial distribution of entry factors (e.g., SR-B1, Claudin-1, and Occludin) present in the polarized human liver (Reynolds *et al*., 2008; Zeisel *et al*, 2011). To bridge this gap, further investigations in biologically relevant systems—including primary human hepatocytes and polarized 3D liver organoids—are warranted. Evaluating mini-19 in these native-like tissue microenvironments will be a critical step prior to systemic *in vivo* validation.

In summary, mini-19 represents a next-generation entry inhibitor that bridges the gap between small molecules and biologics. By combining sub-nanomolar affinity, exceptional biophysical robustness, and a high genetic barrier to resistance, mini-19 establishes a potent alternative to conventional antibody-based therapies. Its clinical utility is maximized when utilized in a combinational framework with existing DAAs. While DAAs suppress intracellular viral replication, they are ineffective against initial attachment and cell-to-cell transmission. By targeting the host-side entry machinery, mini-19 complements existing DAAs by addressing these inherent limitations of DAAs, potentially preventing viral reseeding and the selection of resistance-associated variants. Following advanced *ex vivo* assessments, *in vivo* validation using humanized mouse models will be essential to confirm clinical efficacy and explore broader utility in blocking other CD81-dependent pathogens. By targeting a critical, early step in the HCV life cycle, mini-19 not only complements current DAA strategies but also addresses specific clinical scenarios—such as prophylactic use and the prevention of graft reinfection—where current therapeutic options remain insufficient.

## Data Availability

All data supporting the findings of this study are available within the main paper and its Expanded View (EV) figures and tables. Raw data, structural coordinates, and simulation trajectories are fully prepared and will be provided during the peer review process.

## Author Contributions

S.A. conceived the project, led the computational protein design, and conducted the principal biochemical experiments (including protein purification, SPR, and CD). E.J. performed the HCV neutralization assays. H.S.C. assisted with the computational modeling, SPR analysis, and data interpretation. S.S.K. executed the EV assays and the HEK293 cell binding assays. D.W.K. conducted the human PBMC binding assays, and W.Y. performed dSTORM imaging and analysis. S.K.J., S.W.Y., S.-J.H., Y.S.G., and K.H.P.P. provided technical guidance and conceptual advice. H.-S.C. and S.K. directed the overall research project. S.A. wrote the original manuscript draft. All authors contributed to data interpretation and manuscript preparation, and approved the final version.

## Supplementary Information

Supplementary information accompanies the manuscript on the *The EMBO Journal* website.

---

**Figure EV1:**
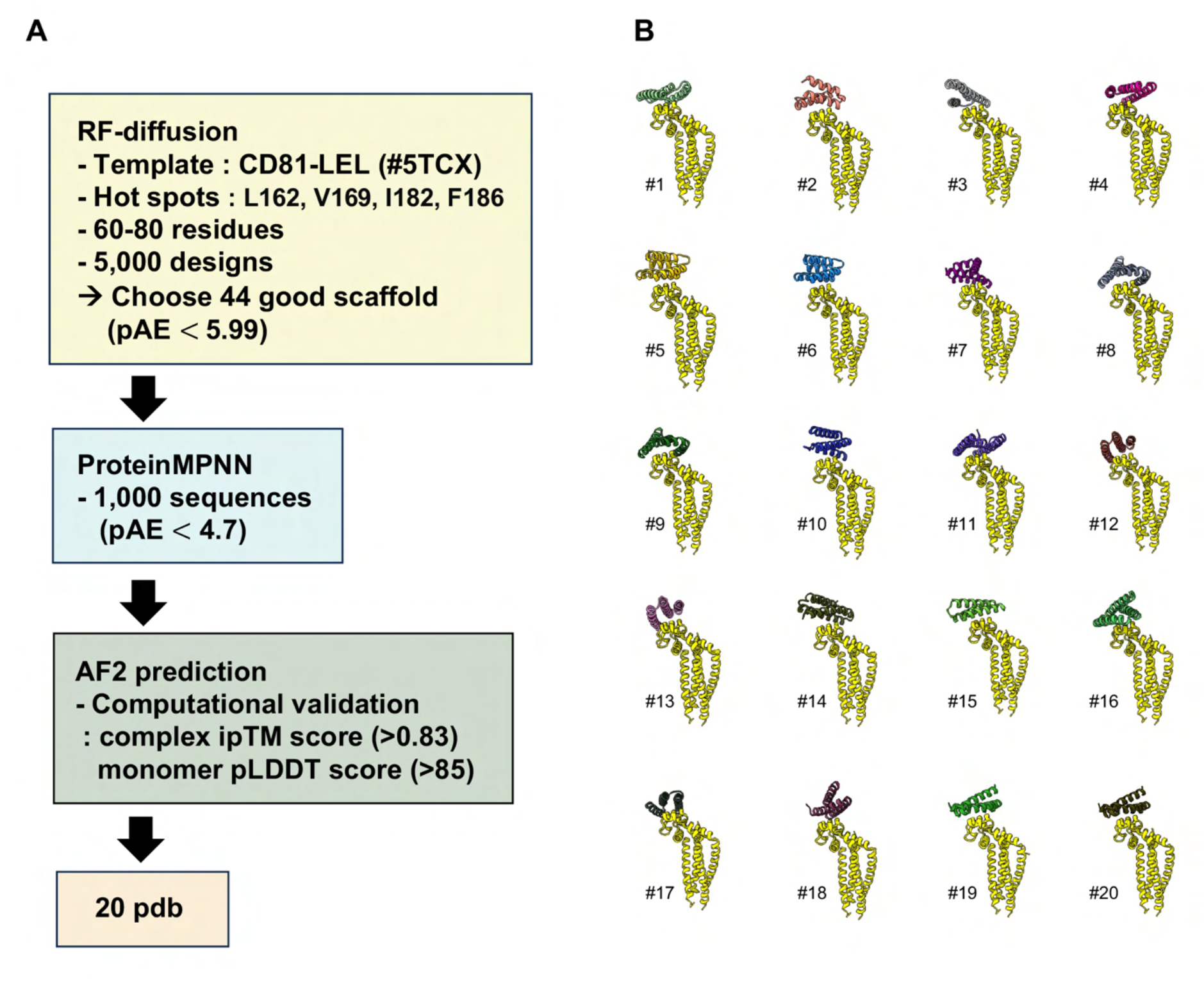
Computational design workflow and structural selection of *de novo* CD81-targeting mini-proteins. **(A)** Schematic representation of the *de novo* design pipeline. High-affinity mini-proteins were engineered through a multi-step computational framework. Initially, RFdiffusion was employed to generate 5,000 unique scaffolds (60–80 amino acids in length), utilizing the human CD81 large extracellular loop (LEL) domain (PDB: 5TCX) as the structural template. From this library, 44 scaffolds were prioritized based on a predicted Alignment Error (pAE) threshold of < 5.99. Subsequent sequence optimization was performed using ProteinMPNN (1,000 sequences per scaffold). The resulting candidates were further filtered using a stricter pAE threshold (< 4.7) and computationally validated using AlphaFold2 (AF2). Final selection required a multimer interface predicted Template Modeling (ipTM) score > 0.83 and a monomer predicted Local Distance Difference Test (pLDDT) score > 85. This pipeline culminated in the selection of 20 high-confidence candidates for experimental characterization. **(B)** Structural models of the 20 selected mini-protein candidates. Ribbon representations illustrate the diverse *de novo* designed mini-proteins (various colors) in complex with the CD81 target (yellow). Each model (#1–#20) highlights the varied binding modes and helical topologies achieved through the diffusion-based design process.

**Figure EV2:**
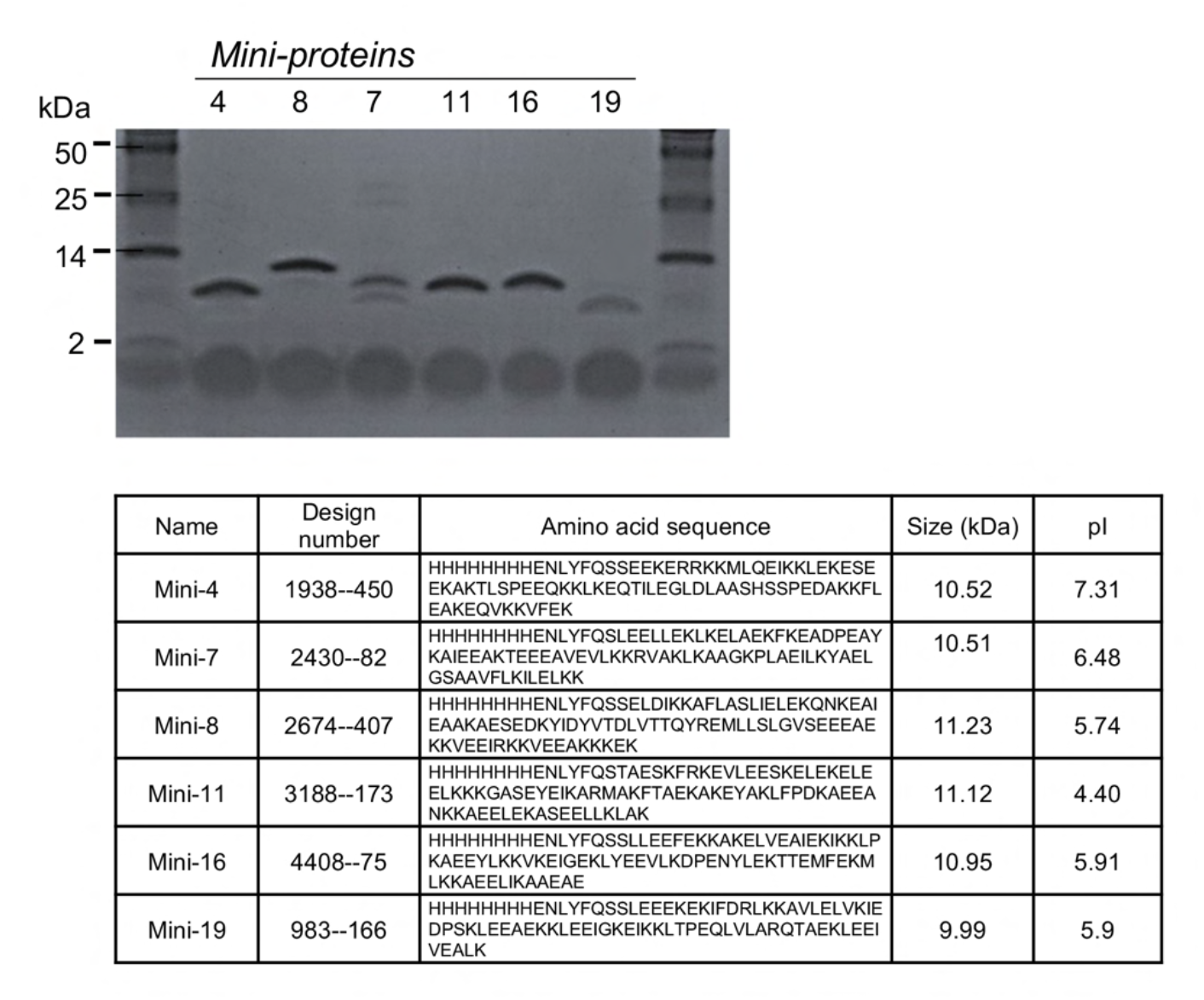
Purification profile and sequence specifications of the CD81-targeting mini-proteins. (Top) Representative Tricine SDS-PAGE gel showing the successful recombinant expression and purification of the six prioritized candidates. While mini-7 displays multiple bands indicative of potential oligomerization or minor impurities, the remaining candidates exhibit sharp, singular bands. (Bottom) Summary table outlining the biophysical properties of the mini-proteins. The table provides unique design numbers, full amino acid sequences (including the N-terminal 8xHis-TEV tag: HHHHHHHHENLYFQS), theoretical molecular weights, and isoelectric points (pI). The migration of the purified mini-proteins on the gel is consistent with their calculated molecular weights and pI.

**Figure EV3:**
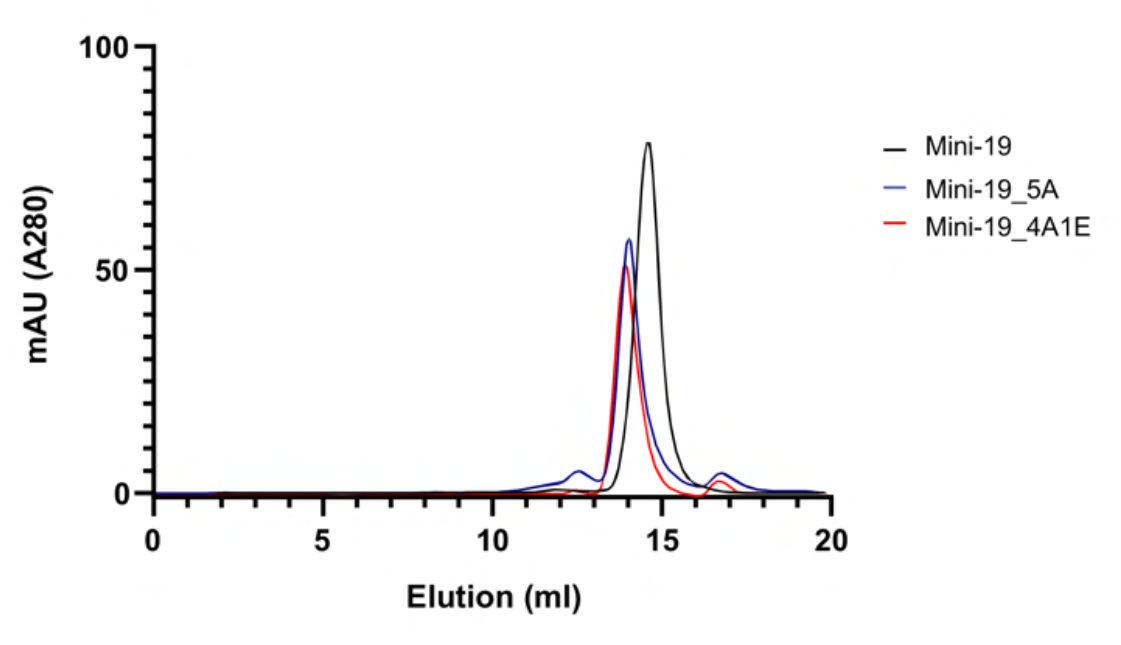
Size-exclusion chromatography profiles of the designed mini-proteins. The purified Mini-19 and its binding-deficient mutants were analyzed using a Superdex 75 Increase 10/300 GL column (Cytiva) equilibrated with 20 mM HEPES (pH 7.5), 150 mM NaCl.

**Figure EV4:**
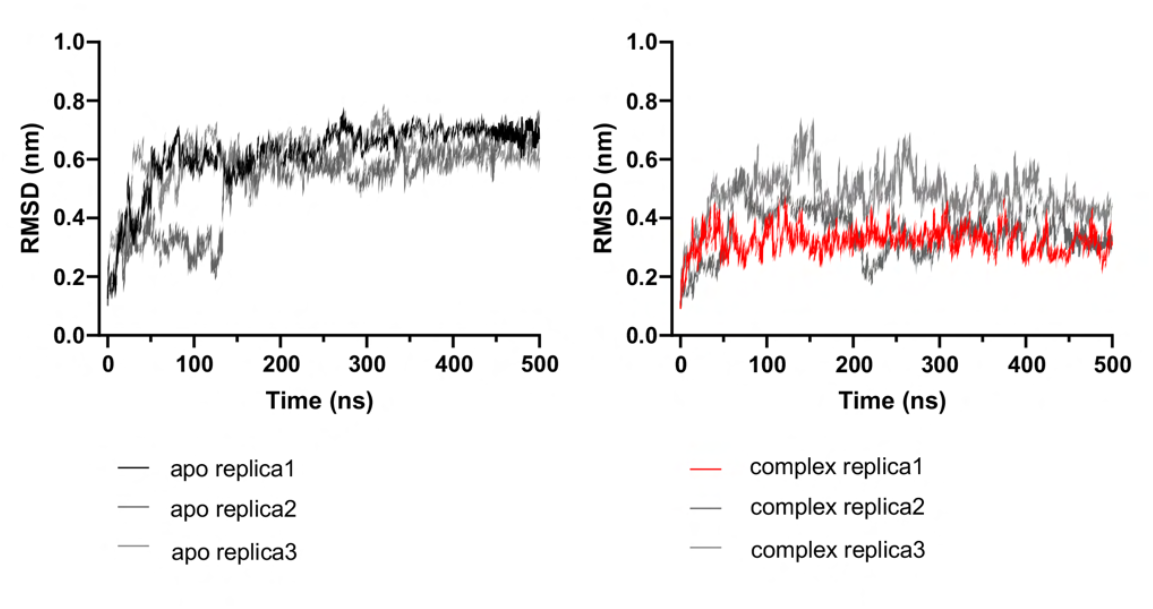
RMSD analysis of the CD81-LEL in the apo and mini-19-bound states. Backbone RMSD trajectories from three independent 500 ns MD simulation replicas for the apo (left) and complex (right) states, performed at an acidic endosomal pH (pH 4.0). The representative trajectories shown in Fig. 2D are highlighted in black (apo) and red (complex).

**Figure EV5:**
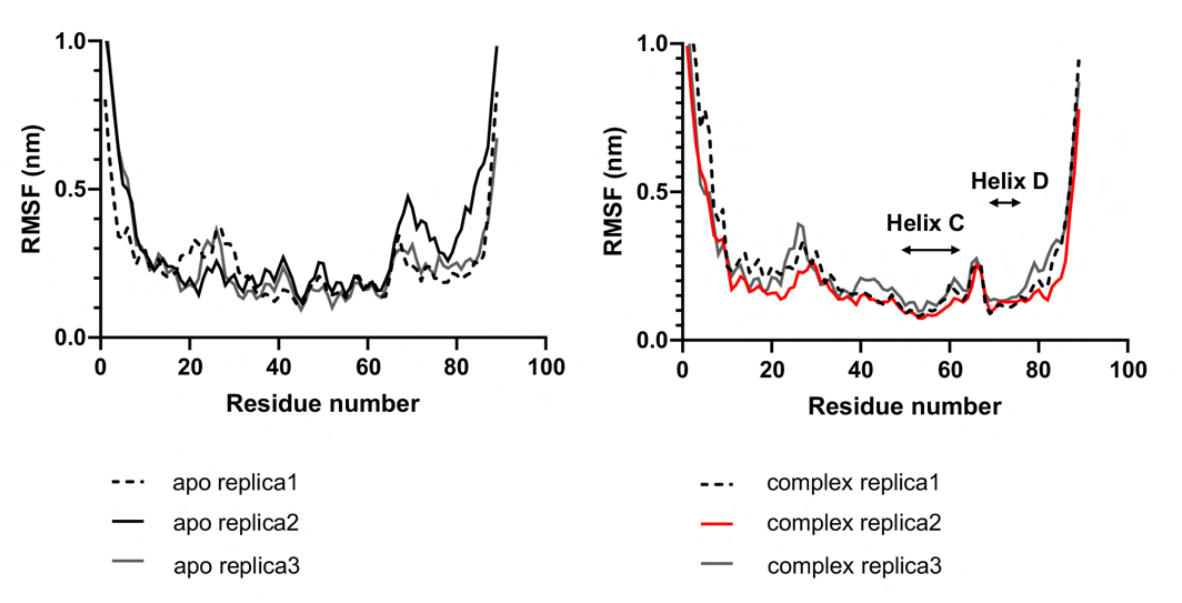
RMSF analysis of the CD81-LEL in the apo and mini-19-bound states. Backbone RMSF profiles from three independent 500 ns MD simulation replicas for the apo (left) and complex (right) states, performed at pH 4.0. The representative profiles shown in Fig. 2E are highlighted as solid black (apo) and red (complex) lines.

**Figure EV6:**
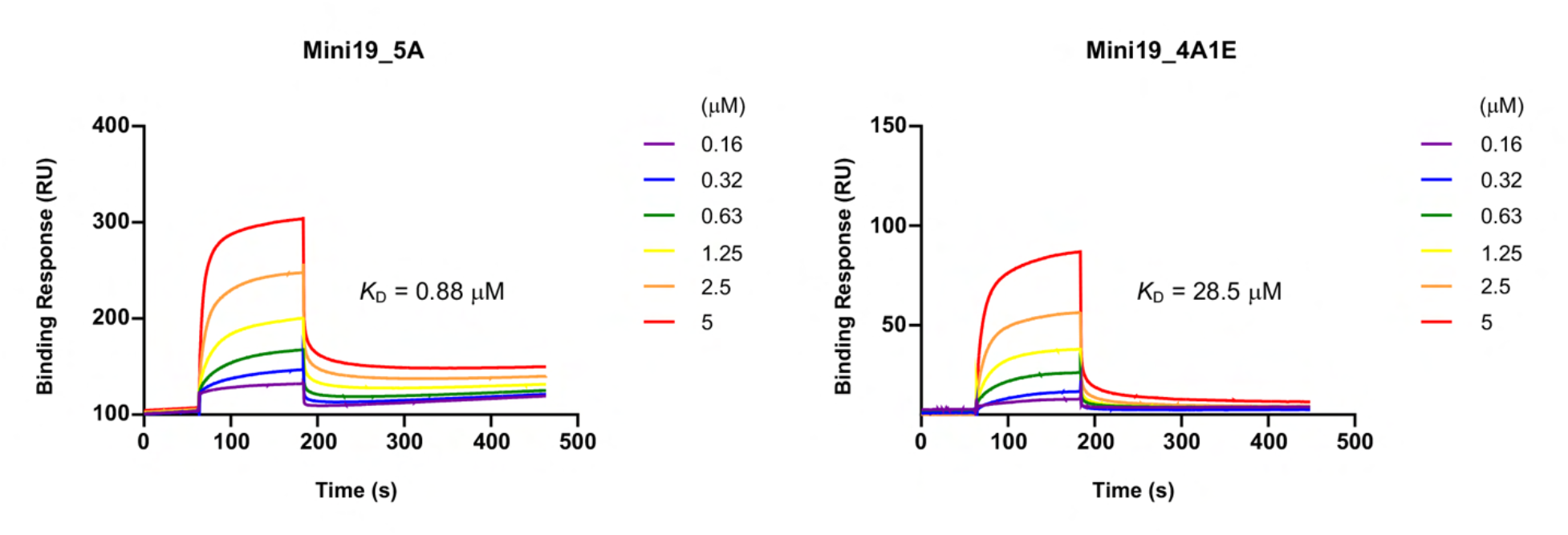
SPR analysis of CD81-binding-deficient mini-19 mutants. SPR sensorgrams and steady-state binding curves are shown for the rationally designed binding-deficient mutants, 5A and 4A1E, to validate the predicted CD81–mini-19 interface. The *y*-axis represents the binding response in resonance units (RU).

**Figure EV7:**
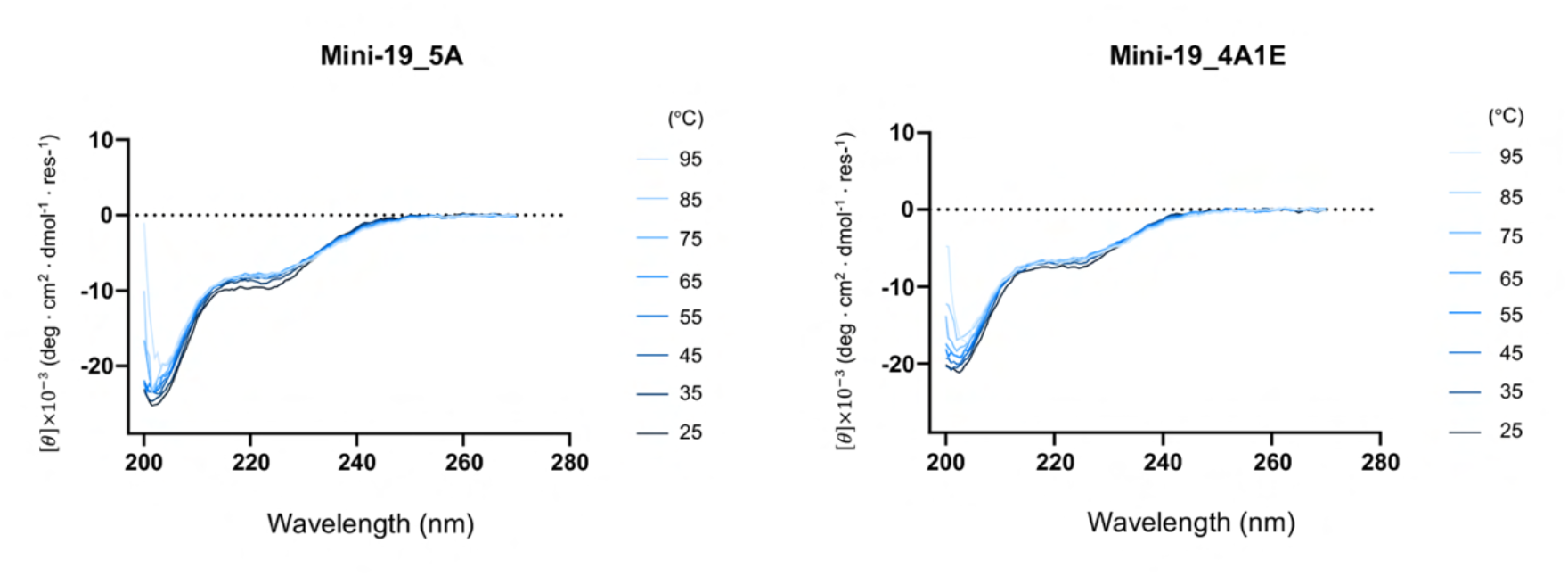
Circular dichroism (CD) analysis of CD81-binding-deficient mini-19 mutants. Far-UV CD spectra of the rationally designed binding-deficient mutants, 5A and 4A1E, were measured across a temperature range of 25°C to 95°C to evaluate their secondary structure and thermal stability. The *y*-axis represents mean residue ellipticity ([θ]).

**Figure EV8:**
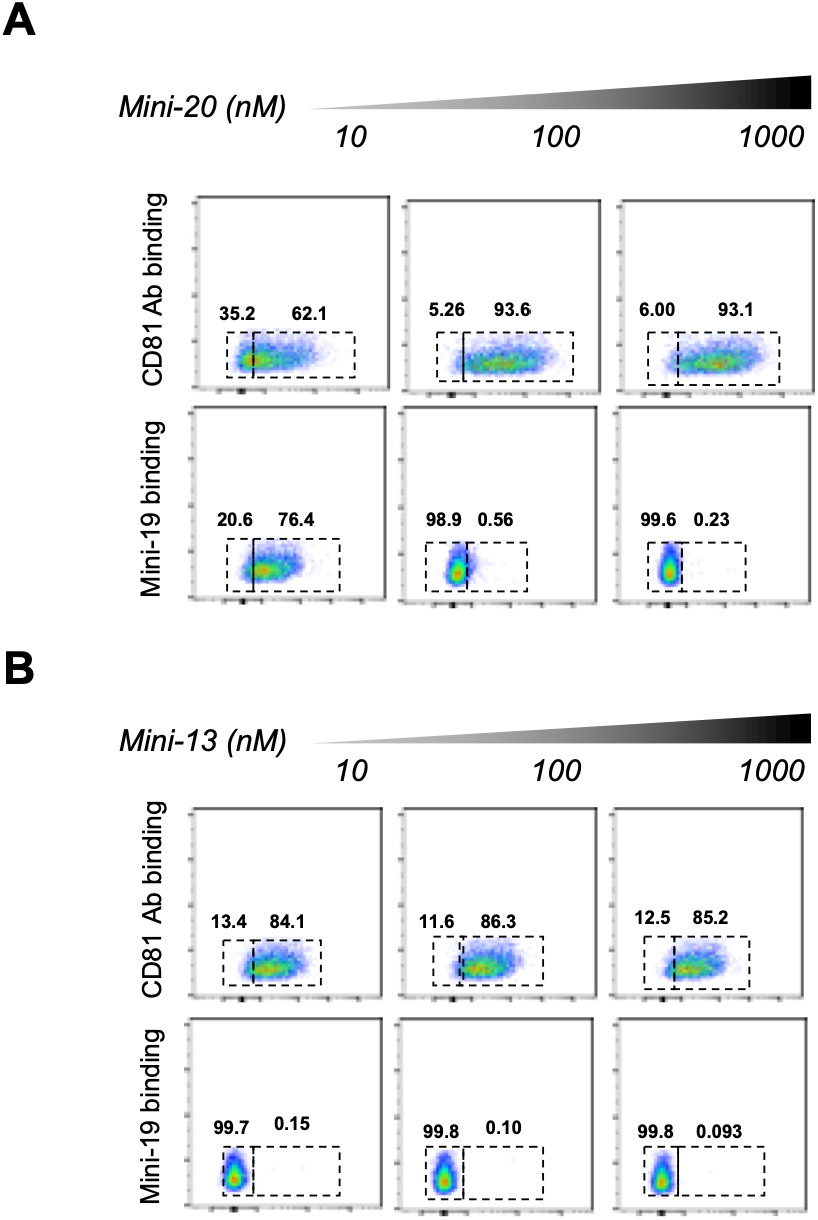
Flow cytometric evaluation of additional *de novo* designed mini-proteins using human PBMCs. (**A**) Binding analysis of mini-20. Mini-20, which shares the same structural scaffold as mini-19 but incorporates sequence variations, demonstrates effective target engagement and competitive displacement of the αCD81 monoclonal antibodies on CD19⁺ B cells within the human PBMC population, exhibiting a similar but slightly attenuated efficacy compared to mini-19. (**B**) Binding analysis of mini-13. Despite possessing a computationally designed binding interface, the negative control mini-13 exhibited neither detectable binding to CD19⁺ B cells nor competitive displacement of the αCD81 monoclonal antibodies, even at elevated concentrations (10 to 1,000 nM).

**Figure EV9:**
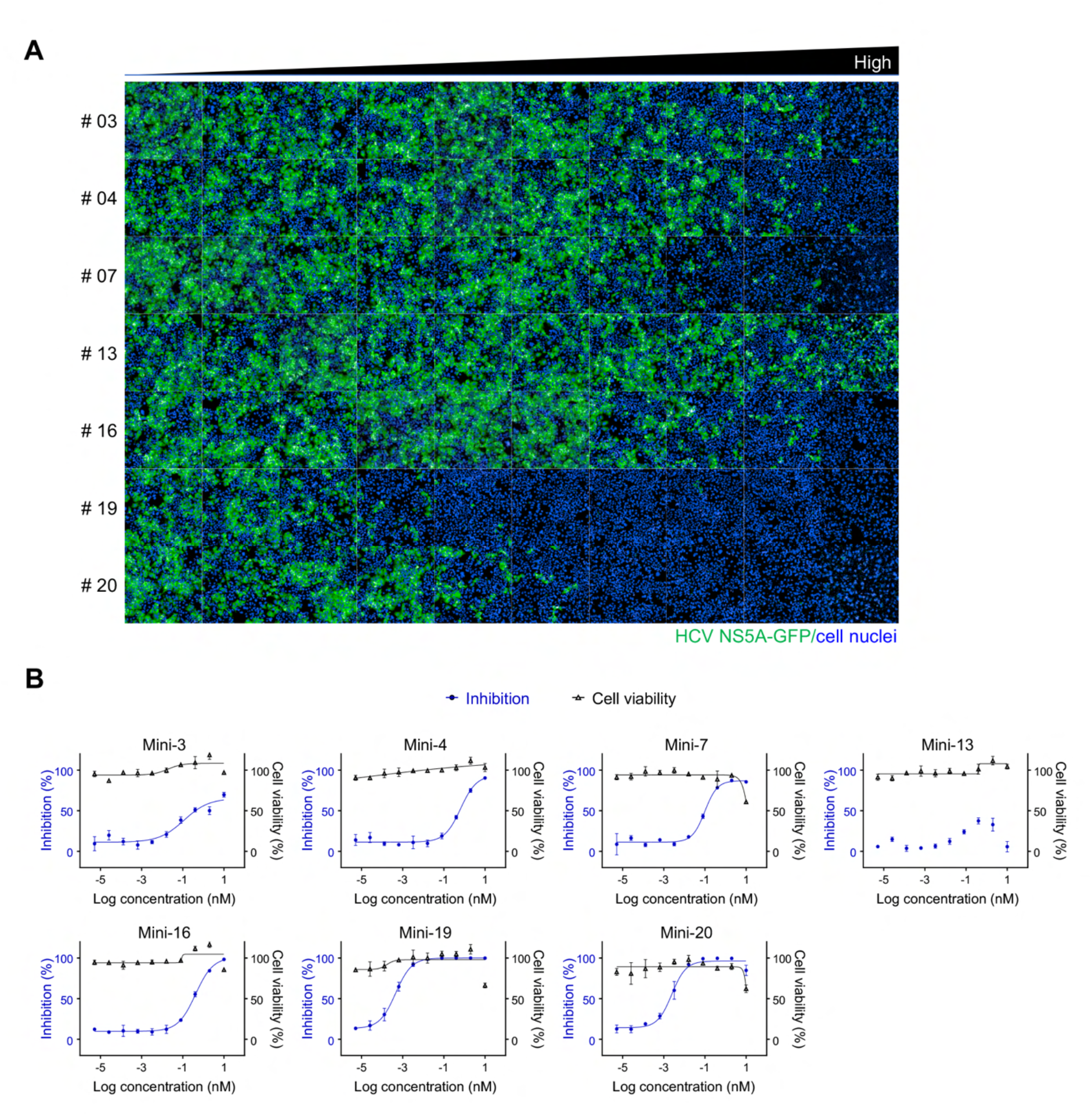
Antiviral efficacy of CD81-targeting mini-proteins against HCVcc infection. **(A)** Representative confocal microscopy images of HCVcc-infected cells treated with CD81-targeting mini-proteins. Cells were treated with six active candidates (mini-03, 04, 07, 16, 19, and 20) and a negative control (mini-13) using a 10-point, 5-fold serial dilution starting at 100 µM. At 3 days post-infection (dpi), cells were fixed and visualized for HCV infection (GFP, green) and cell nuclei (Hoechst 33342, blue). Images represent one of nine subfields acquired per well. **(B)** Dose-response curves depicting the antiviral activity and cytotoxicity of the mini-protein candidates. Percent inhibition (blue circles) and percent cell viability (black triangles) were determined as described in the Materials and Methods.

**Figure EV10:**
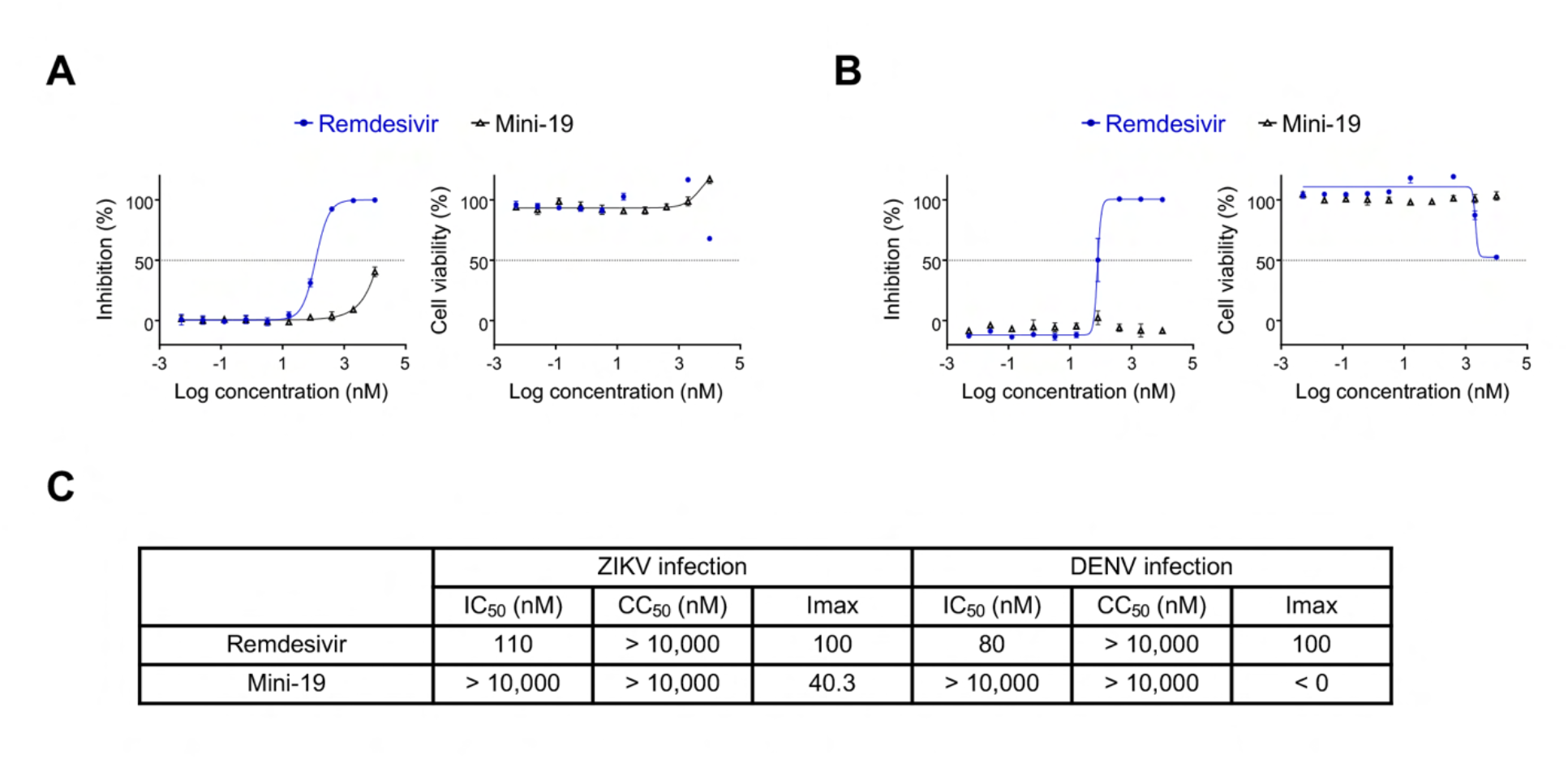
Antiviral activity of mini-19 against ZIKV and DENV. (A and. **B)** Dose-response inhibition of Zika virus (A; African strain, MOI = 0.1) and dengue virus (B; serotype 1, MOI = 0.2) in cells pre-treated with inhibitors for 1 h. Viral infection was quantified at 2 days post-infection (dpi) by an immunofluorescence assay using an αE protein antibody (4G2). Mini-19 (black triangles) was tested using a 10-point, 5-fold serial dilution starting from 10 µM. Remdesivir (blue circles) served as a reference inhibitor. All conditions were tested in triplicate. Data were normalized using 3 µM remdesivir as the 100% inhibition control and untreated cells as the 0% inhibition control. **(C)** Summary of antiviral efficacy. Activity is presented as IC50, CC50, and maximum inhibition (Imax) values.

**Table EV1:**
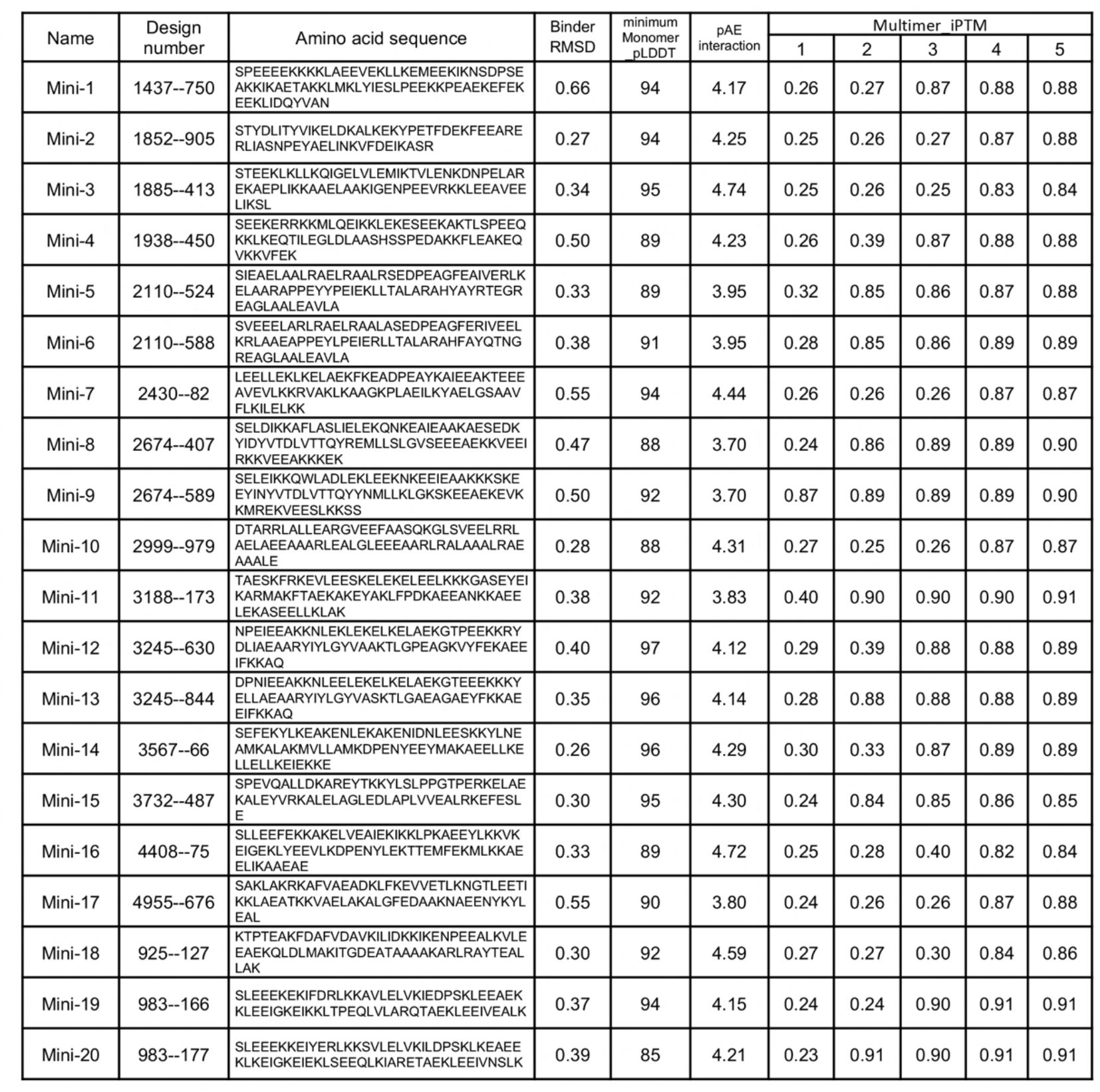
*In silico* evaluation metrics and sequences of *de novo* designed CD81-targeting mini-proteins. This table summarizes the computational validation metrics and sequences for the top 20 candidate mini-proteins generated from the design pipeline described in Figure EV1. To rigorously evaluate binding confidence and eliminate false positives—such as mis-targeted binding to exposed hydrophobic surfaces of the truncated CD81 template—AlphaFold2 (AF2) multimer predictions were utilized. The minimum Monomer pLDDT reflects the predicted folding confidence of the isolated scaffold (threshold > 85). Interaction accuracy was assessed using predicted Aligned Error at the interface (pAE interaction, threshold < 4.7). The Multimer_ipTM columns display the interface predicted Template Modeling scores across five independent AF2 prediction models. To ensure high target specificity and structural convergence at the intended LEL hotspot, candidates were advanced for *in vitro* validation only if at least two of the five independent AF2 models achieved an ipTM score > 0.83 with identical backbone conformations.

**Table EV2:** Binding affinities of selected CD81-targeting mini-proteins determined by SPR. Equilibrium dissociation constants (*K*D) are presented for eight representative mini-protein candidates. While several initial designs exhibited affinities in the nanomolar range (50–382 nM), mini-19 emerged as the most potent lead with an exceptional sub-nanomolar affinity of 0.5 nM.

| # | $K_D$ |
| --- | --- |
| Mini-3 | 201 nM |
| Mini-4 | 260 nM |
| Mini-7 | 50 nM |
| Mini-8 | 382 nM |
| Mini-11 | 203 nM |
| Mini-16 | 117 nM |
| **Mini-19** | **0.5 nM** |
| Mini-20 | 24 nM |

## References

Ahmed W, Neelakanta G, Sultana H (2021) Tetraspanins as Potential Therapeutic Candidates for Targeting Flaviviruses. Front Immunol 12: 630571

Al Olaby RR, Cocquerel L, Zemla A, Saas L, Dubuisson J, Vielmetter J, Marcotrigiano J, Khan AG, Vences Catalan F, Perryman AL et al (2014) Identification of a novel drug lead that inhibits HCV infection and cell-to-cell transmission by targeting the HCV E2 glycoprotein. PLoS One 9: e111333

Aljowaie RM, Almajhdi FN, Ali HH, El-Wetidy MS, Shier MK (2020) Inhibition of hepatitis C virus genotype 4 replication using siRNA targeted to the viral core region and the CD81 cellular receptor. Cell Stress Chaperones 25: 345–355

Bailly C, Thuru X (2023) Targeting of Tetraspanin CD81 with Monoclonal Antibodies and Small Molecules to Combat Cancers and Viral Diseases. Cancers (Basel) 15

Berger S, Seeger F, Yu TY, Aydin M, Yang H, Rosenblum D, Guenin-Mace L, Glassman C, Arguinchona L, Sniezek C et al (2024) Preclinical proof of principle for orally delivered Th17 antagonist miniproteins. Cell 187: 4305–4317 e4318

Blight KJ, McKeating JA, Rice CM (2002) Highly permissive cell lines for subgenomic and genomic hepatitis C virus RNA replication. J Virol 76: 13001–13014

Brazzoli M, Bianchi A, Filippini S, Weiner A, Zhu Q, Pizza M, Crotta S (2008) CD81 is a central regulator of cellular events required for hepatitis C virus infection of human hepatocytes. J Virol 82: 8316–8329

Brown RS (2005) Hepatitis C and liver transplantation. Nature 436: 973–978

Bush CO, Pokrovskii MV, Saito R, Morganelli P, Canales E, Clarke MO, Lazerwith SE, Golde J, Reid BG, Babaoglu K et al (2014) A small-molecule inhibitor of hepatitis C virus infectivity. Antimicrob Agents Chemother 58: 386–396

Cao L, Coventry B, Goreshnik I, Huang B, Sheffler W, Park JS, Jude KM, Markovic I, Kadam RU, Verschueren KHG et al (2022) Design of protein-binding proteins from the target structure alone. Nature 605: 551–560

Cao L, Goreshnik I, Coventry B, Case JB, Miller L, Kozodoy L, Chen RE, Carter L, Walls AC, Park YJ et al (2020) De novo design of picomolar SARS-CoV-2 miniprotein inhibitors. Science 370: 426–431

Carloni V, Mazzocca A, Ravichandran KS (2004) Tetraspanin CD81 is linked to ERK/MAPKinase signaling by Shc in liver tumor cells. Oncogene 23: 1566–1574

Cherukuri A, Shoham T, Sohn HW, Levy S, Brooks S, Carter R, Pierce SK (2004) The tetraspanin CD81 is necessary for partitioning of coligated CD19/CD21-B cell antigen receptor complexes into signaling-active lipid rafts. J Immunol 172: 370–380

Choi D, Go G, Kim DK, Lee J, Park SM, Di Vizio D, Gho YS (2020) Quantitative proteomic analysis of trypsin-treated extracellular vesicles to identify the real-vesicular proteins. J Extracell Vesicles 9: 1757209

Cunha ES, Sfriso P, Rojas AL, Roversi P, Hospital A, Orozco M, Abrescia NGA (2017) Mechanism of Structural Tuning of the Hepatitis C Virus Human Cellular Receptor CD81 Large Extracellular Loop. Structure 25: 53–65

Cuypers L, Ceccherini-Silberstein F, Van Laethem K, Li G, Vandamme AM, Rockstroh JK (2016) Impact of HCV genotype on treatment regimens and drug resistance: a snapshot in time. Rev Med Virol 26: 408–434

Das D, Pandya M (2018) Recent Advancement of Direct-acting Antiviral Agents (DAAs) in Hepatitis C Therapy. Mini Rev Med Chem 18: 584–596

Dauparas J, Anishchenko I, Bennett N, Bai H, Ragotte RJ, Milles LF, Wicky BIM, Courbet A, de Haas RJ, Bethel N et al (2022) Robust deep learning-based protein sequence design using ProteinMPNN. Science 378: 49–56

Desombere I, Mesalam AA, Urbanowicz RA, Van Houtte F, Verhoye L, Keck ZY, Farhoudi A, Vercauteren K, Weening KE, Baumert TF et al (2017) A novel neutralizing human monoclonal antibody broadly abrogates hepatitis C virus infection in vitro and in vivo. Antiviral Res 148: 53–64

Ding Q, von Schaewen M, Hrebikova G, Heller B, Sandmann L, Plaas M, Ploss A (2017) Mice Expressing Minimally Humanized CD81 and Occludin Genes Support Hepatitis C Virus Uptake In Vivo. J Virol 91

Douam F, Lavillette D, Cosset FL (2015) The mechanism of HCV entry into host cells. Prog Mol Biol Transl Sci 129: 63–107

Drummer HE, Wilson KA, Poumbourios P (2002) Identification of the hepatitis C virus E2 glycoprotein binding site on the large extracellular loop of CD81. J Virol 76: 11143–11147

Evans MJ, von Hahn T, Tscherne DM, Syder AJ, Panis M, Wolk B, Hatziioannou T, McKeating JA, Bieniasz PD, Rice CM (2007) Claudin-1 is a hepatitis C virus co-receptor required for a late step in entry. Nature 446: 801–805

Fasano M, Ieva F, Ciarallo M, Caccianotti B, Santantonio TA (2024) Acute Hepatitis C: Current Status and Future Perspectives. Viruses 16

Feneant L, Levy S, Cocquerel L (2014) CD81 and hepatitis C virus (HCV) infection. Viruses 6: 535–572

Florin L, Lang T (2018) Tetraspanin Assemblies in Virus Infection. Front Immunol 9: 1140

Fofana I, Xiao F, Thumann C, Turek M, Zona L, Tawar RG, Grunert F, Thompson J, Zeisel MB, Baumert TF (2013) A novel monoclonal anti-CD81 antibody produced by genetic immunization efficiently inhibits Hepatitis C virus cell-cell transmission. PLoS One 8: e64221

Gerold G, Moeller R, Pietschmann T (2020) Hepatitis C Virus Entry: Protein Interactions and Fusion Determinants Governing Productive Hepatocyte Invasion. Cold Spring Harb Perspect Med 10

Grakoui A, McCourt DW, Wychowski C, Feinstone SM, Rice CM (1993a) Characterization of the hepatitis C virus-encoded serine proteinase: determination of proteinase-dependent polyprotein cleavage sites. J Virol 67: 2832–2843

Grakoui A, McCourt DW, Wychowski C, Feinstone SM, Rice CM (1993b) A second hepatitis C virus-encoded proteinase. Proc Natl Acad Sci U S A 90: 10583–10587

Greenfield NJ (2006) Using circular dichroism collected as a function of temperature to determine the thermodynamics of protein unfolding and binding interactions. Nat Protoc 1: 2527–2535

Hammerstad SS, Stefan M, Blackard J, Owen RP, Lee HJ, Concepcion E, Yi Z, Zhang W, Tomer Y (2017) Hepatitis C Virus E2 Protein Induces Upregulation of IL-8 Pathways and Production of Heat Shock Proteins in Human Thyroid Cells. J Clin Endocrinol Metab 102: 689–697

Harris HJ, Davis C, Mullins JG, Hu K, Goodall M, Farquhar MJ, Mee CJ, McCaffrey K, Young S, Drummer H et al (2010) Claudin association with CD81 defines hepatitis C virus entry. J Biol Chem 285: 21092–21102

Harris HJ, Farquhar MJ, Mee CJ, Davis C, Reynolds GM, Jennings A, Hu K, Yuan F, Deng H, Hubscher SG et al (2008) CD81 and claudin 1 coreceptor association: role in hepatitis C virus entry. J Virol 82: 5007–5020

Hemler ME (2005) Tetraspanin functions and associated microdomains. Nat Rev Mol Cell Biol 6: 801–811

Higginbottom A, Quinn ER, Kuo CC, Flint M, Wilson LH, Bianchi E, Nicosia A, Monk PN, McKeating JA, Levy S (2000) Identification of amino acid residues in CD81 critical for interaction with hepatitis C virus envelope glycoprotein E2. J Virol 74: 3642–3649

Houldsworth A, Metzner MM, Demaine A, Hodgkinson A, Kaminski E, Cramp M (2014) CD81 sequence and susceptibility to hepatitis C infection. J Med Virol 86: 162–168

Huang B, Coventry B, Borowska MT, Arhontoulis DC, Exposit M, Abedi M, Jude KM, Halabiya SF, Allen A, Cordray C et al (2024) De novo design of miniprotein antagonists of cytokine storm inducers. Nat Commun 15: 7064

Ji C, Liu Y, Pamulapati C, Bohini S, Fertig G, Schraeml M, Rubas W, Brandt M, Ries S, Ma H et al (2015) Prevention of hepatitis C virus infection and spread in human liver chimeric mice by an anti-CD81 monoclonal antibody. Hepatology 61: 1136–1144

Kanwal F, Kramer JR, Asch SM, Cao Y, Li L, El-Serag HB (2020) Long-Term Risk of Hepatocellular Carcinoma in HCV Patients Treated With Direct Acting Antiviral Agents. Hepatology 71: 44–55

Kim OY, Park HT, Dinh NTH, Choi SJ, Lee J, Kim JH, Lee SW, Gho YS (2017) Bacterial outer membrane vesicles suppress tumor by interferon-gamma-mediated antitumor response. Nat Commun 8: 626

Kitadokoro K, Bordo D, Galli G, Petracca R, Falugi F, Abrignani S, Grandi G, Bolognesi M (2001) CD81 extracellular domain 3D structure: insight into the tetraspanin superfamily structural motifs. EMBO J 20: 12–18

Koutsoudakis G, Herrmann E, Kallis S, Bartenschlager R, Pietschmann T (2007) The level of CD81 cell surface expression is a key determinant for productive entry of hepatitis C virus into host cells. J Virol 81: 588–598

Krieger SE, Zeisel MB, Davis C, Thumann C, Harris HJ, Schnober EK, Mee C, Soulier E, Royer C, Lambotin M et al (2010) Inhibition of hepatitis C virus infection by anti-claudin-1 antibodies is mediated by neutralization of E2-CD81-claudin-1 associations. Hepatology 51: 1144–1157

Kumar A, Hossain RA, Yost SA, Bu W, Wang Y, Dearborn AD, Grakoui A, Cohen JI, Marcotrigiano J (2021) Structural insights into hepatitis C virus receptor binding and entry. Nature 598: 521–525

Lee M, Yang J, Jo E, Lee JY, Kim HY, Bartenschlager R, Shin EC, Bae YS, Windisch MP (2017) A Novel Inhibitor IDPP Interferes with Entry and Egress of HCV by Targeting Glycoprotein E1 in a Genotype-Specific Manner. Sci Rep 7: 44676

Levy S, Shoham T (2005) The tetraspanin web modulates immune-signalling complexes. Nat Rev Immunol 5: 136–148

Lin LT, Chung CY, Hsu WC, Chang SP, Hung TC, Shields J, Russell RS, Lin CC, Li CF, Yen MH et al (2015) Saikosaponin b2 is a naturally occurring terpenoid that efficiently inhibits hepatitis C virus entry. J Hepatol 62: 541–548

Maecker HT, Levy S (1997) Normal lymphocyte development but delayed humoral immune response in CD81-null mice. J Exp Med 185: 1505–1510

Martin F, Roth DM, Jans DA, Pouton CW, Partridge LJ, Monk PN, Moseley GW (2005) Tetraspanins in viral infections: a fundamental role in viral biology? J Virol 79: 10839–10851

Masciopinto F, Giovani C, Campagnoli S, Galli-Stampino L, Colombatto P, Brunetto M, Yen TS, Houghton M, Pileri P, Abrignani S (2004) Association of hepatitis C virus envelope proteins with exosomes. Eur J Immunol 34: 2834–2842

Meola A, Sbardellati A, Bruni Ercole B, Cerretani M, Pezzanera M, Ceccacci A, Vitelli A, Levy S, Nicosia A, Traboni C et al (2000) Binding of hepatitis C virus E2 glycoprotein to CD81 does not correlate with species permissiveness to infection. J Virol 74: 5933–5938

Meuleman P, Hesselgesser J, Paulson M, Vanwolleghem T, Desombere I, Reiser H, Leroux-Roels G (2008) Anti-CD81 antibodies can prevent a hepatitis C virus infection in vivo. Hepatology 48: 1761–1768

Miyazaki T, Muller U, Campbell KS (1997) Normal development but differentially altered proliferative responses of lymphocytes in mice lacking CD81. EMBO J 16: 4217–4225

Monk PN, Partridge LJ (2012) Tetraspanins: gateways for infection. Infect Disord Drug Targets 12: 4–17

Murakami E, Tolstykh T, Bao H, Niu C, Steuer HM, Bao D, Chang W, Espiritu C, Bansal S, Lam AM et al (2010) Mechanism of activation of PSI-7851 and its diastereoisomer PSI-7977. J Biol Chem 285: 34337–34347

Pan Q, Xie Y, Zhang Y, Guo X, Wang J, Liu M, Zhang XL (2024) EGFR core fucosylation, induced by hepatitis C virus, promotes TRIM40-mediated-RIG-I ubiquitination and suppresses interferon-I antiviral defenses. Nat Commun 15: 652

Perez-Hernandez D, Gutierrez-Vazquez C, Jorge I, Lopez-Martin S, Ursa A, Sanchez-Madrid F, Vazquez J, Yanez-Mo M (2013) The intracellular interactome of tetraspanin-enriched microdomains reveals their function as sorting machineries toward exosomes. J Biol Chem 288: 11649–11661

Pettersen EF, Goddard TD, Huang CC, Meng EC, Couch GS, Croll TI, Morris JH, Ferrin TE (2021) UCSF ChimeraX: Structure visualization for researchers, educators, and developers. Protein Sci 30: 70–82

Pileri P, Uematsu Y, Campagnoli S, Galli G, Falugi F, Petracca R, Weiner AJ, Houghton M, Rosa D, Grandi G et al (1998) Binding of hepatitis C virus to CD81. Science 282: 938–941

Ploss A, Evans MJ, Gaysinskaya VA, Panis M, You H, de Jong YP, Rice CM (2009) Human occludin is a hepatitis C virus entry factor required for infection of mouse cells. Nature 457: 882–886

Ragotte RJ, Liang H, Tam J, Miletic S, Berman JM, Palou R, Weidle C, Li Z, Glogl M, Beilhartz GL et al (2025a) De novo design of potent inhibitors of clostridial family toxins. Proc Natl Acad Sci U S A 122: e2509329122

Ragotte RJ, Tortorici MA, Catanzaro NJ, Addetia A, Coventry B, Froggatt HM, Lee J, Stewart C, Brown JT, Goreshnik I et al (2025b) Designed miniproteins potently inhibit and protect against MERS-CoV. Cell Rep 44: 115760

Rajesh S, Sridhar P, Tews BA, Feneant L, Cocquerel L, Ward DG, Berditchevski F, Overduin M (2012) Structural basis of ligand interactions of the large extracellular domain of tetraspanin CD81. J Virol 86: 9606–9616

Ramakrishnaiah V, Thumann C, Fofana I, Habersetzer F, Pan Q, de Ruiter PE, Willemsen R, Demmers JA, Stalin Raj V, Jenster G et al (2013) Exosome-mediated transmission of hepatitis C virus between human hepatoma Huh7.5 cells. Proc Natl Acad Sci U S A 110: 13109–13113

Reynolds GM, Harris HJ, Jennings A, Hu K, Grove J, Lalor PF, Adams DH, Balfe P, Hubscher SG, McKeating JA (2008) Hepatitis C virus receptor expression in normal and diseased liver tissue. Hepatology 47: 418–427

Rice JP (2015) Hepatitis C treatment: Back to the warehouse. Clin Liver Dis (Hoboken) 6: 27–29

Risueno C, Carbajo I, Charro D, Abrescia NGA, Coluzza I (2025) pH-antenna residues trigger a large-scale conformational change in the large extracellular loop domain of the CD81 human receptor. J Chem Phys 162

Scarselli E, Ansuini H, Cerino R, Roccasecca RM, Acali S, Filocamo G, Traboni C, Nicosia A, Cortese R, Vitelli A (2002) The human scavenger receptor class B type I is a novel candidate receptor for the hepatitis C virus. EMBO J 21: 5017–5025

Schagger H, von Jagow G (1987) Tricine-sodium dodecyl sulfate-polyacrylamide gel electrophoresis for the separation of proteins in the range from 1 to 100 kDa. Anal Biochem 166: 368–379

Shahid I, Alzahrani AR, Al-Ghamdi SS, Alanazi IM, Rehman S, Hassan S (2021) Hepatitis C Diagnosis: Simplified Solutions, Predictive Barriers, and Future Promises. Diagnostics (Basel) 11

Sharma NR, Mateu G, Dreux M, Grakoui A, Cosset FL, Melikyan GB (2011) Hepatitis C virus is primed by CD81 protein for low pH-dependent fusion. J Biol Chem 286: 30361–30376

Shi Y, Du L, Lv D, Li Y, Zhang Z, Huang X, Tang H (2021) Emerging role and therapeutic application of exosome in hepatitis virus infection and associated diseases. J Gastroenterol 56: 336–349

Silvie O, Rubinstein E, Franetich JF, Prenant M, Belnoue E, Renia L, Hannoun L, Eling W, Levy S, Boucheix C et al (2003) Hepatocyte CD81 is required for Plasmodium falciparum and Plasmodium yoelii sporozoite infectivity. Nat Med 9: 93–96

Smith DA, Fernandez-Antunez C, Magri A, Bowden R, Chaturvedi N, Fellay J, McLauchlan J, Foster GR, Irving WL, Consortium S-H et al (2021) Viral genome wide association study identifies novel hepatitis C virus polymorphisms associated with sofosbuvir treatment failure. Nat Commun 12: 6105

Susa KJ, Rawson S, Kruse AC, Blacklow SC (2021) Cryo-EM structure of the B cell co-receptor CD19 bound to the tetraspanin CD81. Science 371: 300–305

Susa KJ, Seegar TC, Blacklow SC, Kruse AC (2020) A dynamic interaction between CD19 and the tetraspanin CD81 controls B cell co-receptor trafficking. Elife 9

Tsitsikov EN, Gutierrez-Ramos JC, Geha RS (1997) Impaired CD19 expression and signaling, enhanced antibody response to type II T independent antigen and reduction of B-1 cells in CD81-deficient mice. Proc Natl Acad Sci U S A 94: 10844–10849

Vazquez Torres S, Benard Valle M, Mackessy SP, Menzies SK, Casewell NR, Ahmadi S, Burlet NJ, Muratspahic E, Sappington I, Overath MD et al (2025) De novo designed proteins neutralize lethal snake venom toxins. Nature 639: 225–231

Vences-Catalan F, Kuo CC, Rajapaksa R, Duault C, Andor N, Czerwinski DK, Levy R, Levy S (2019) CD81 is a novel immunotherapeutic target for B cell lymphoma. J Exp Med 216: 1497–1508

Watson JL, Juergens D, Bennett NR, Trippe BL, Yim J, Eisenach HE, Ahern W, Borst AJ, Ragotte RJ, Milles LF et al (2023) De novo design of protein structure and function with RFdiffusion. Nature 620: 1089–1100

World Health Organization, 2025. World Health Organization. Geneva.

Zeisel MB, Fofana I, Fafi-Kremer S, Baumert TF (2011) Hepatitis C virus entry into hepatocytes: molecular mechanisms and targets for antiviral therapies. J Hepatol 54: 566–576

Zhang J, Randall G, Higginbottom A, Monk P, Rice CM, McKeating JA (2004) CD81 is required for hepatitis C virus glycoprotein-mediated viral infection. J Virol 78: 1448–1455

Zhao Q, He K, Zhang X, Xu M, Zhang X, Li H (2022a) Production and immunogenicity of different prophylactic vaccines for hepatitis C virus (Review). Exp Ther Med 24: 474

Zhao Z, Chu M, Guo Y, Yang S, Abudurusuli G, Frutos R, Chen T (2022b) Feasibility of Hepatitis C Elimination in China: From Epidemiology, Natural History, and Intervention Perspectives. Front Microbiol 13: 884598

Zhou H, Yan ZH, Yuan Y, Xing C, Jiang N (2021) The Role of Exosomes in Viral Hepatitis and Its Associated Liver Diseases. Front Med (Lausanne) 8: 782485

Zimmerman B, Kelly B, McMillan BJ, Seegar TCM, Dror RO, Kruse AC, Blacklow SC (2016) Crystal Structure of a Full-Length Human Tetraspanin Reveals a Cholesterol-Binding Pocket. Cell 167: 1041–1051 e1011

Zona L, Tawar RG, Zeisel MB, Xiao F, Schuster C, Lupberger J, Baumert TF (2014) CD81-receptor associations--impact for hepatitis C virus entry and antiviral therapies. Viruses 6: 875–892

